# Confronting false discoveries in single-cell differential expression

**DOI:** 10.1101/2021.03.12.435024

**Authors:** Jordan W. Squair, Matthieu Gautier, Claudia Kathe, Mark A. Anderson, Nicholas D. James, Thomas H. Hutson, Rémi Hudelle, Taha Qaiser, Kaya J. E. Matson, Quentin Barraud, Ariel J. Levine, Gioele La Manno, Michael A. Skinnider, Grégoire Courtine

## Abstract

Differential expression analysis in single-cell transcriptomics enables the dissection of cell-type-specific responses to perturbations such as disease, trauma, or experimental manipulation. While many statistical methods are available to identify differentially expressed genes, the principles that distinguish these methods and their performance remain unclear. Here, we show that the relative performance of these methods is contingent on their ability to account for variation between biological replicates. Methods that ignore this inevitable variation are biased and prone to false discoveries. Indeed, the most widely used methods can discover hundreds of differentially expressed genes in the absence of biological differences. Our results suggest an urgent need for a paradigm shift in the methods used to perform differential expression analysis in single-cell data.

---

The abundance of RNA species informs on the past, present and future state of cells and tissues. By enabling the complete quantification of mRNA populations, RNA sequencing (RNA-seq) has provided unprecedented access to the molecular processes active in a biological sample^1^. Diseases, traumas, and experimental manipulations perturb these processes, which leads to changes in the expression of specific mRNAs. Historically, these altered mRNAs were identified using bulk RNA-seq in non-perturbed versus perturbed tissues^2^. However, biological tissues are composed of multiple cell types, whose responses to a perturbation can differ dramatically. Changes in mRNA abundance within multicellular tissues are confounded by different responses across cell types and changes in the relative abundance of these cell types^3^. Consequently, the resolution of bulk RNA-seq is insufficient to characterize the multifaceted responses to biological perturbations.

Single-cell RNA-seq (scRNA-seq) enables the quantification of RNA abundance at the resolution of individual cells4. The maturation of single-cell technologies now enables large-scale comparisons of cell states within complex tissues, thus providing the appropriate resolution to dissect cell-type-specific responses to perturbation^5,6^. The sparsity and heterogeneity of single-cell data initially encouraged the development of specialized statistical methods to identify differentially expressed mRNAs^7,8^. The proliferation of statistical methods for differential expression analysis prompted investigators to ask which methods produced the most biologically accurate results. To answer this question, investigators turned to simulations in an attempt to create a ground truth against which the various methods could be benchmarked. However, simulations require specifying a model from which synthetic patterns of differential expression are generated. Differences in the specification of this model have led investigators to contrasting conclusions^9,10^.

These divergences emphasize the importance of developing a sound epistemological foundation for differential expression in single-cell data^11^. We reasoned that developing such a foundation would require quantifying the performance of the available methods across multiple datasets in which an experimental ground truth is known, and defining the principles that are responsible for differences in performance. We therefore first established a methodological framework that enabled us to curate a resource of ground-truth datasets. Using this resource, we conducted a definitive comparison of the various available methods for differential expression analysis. We found that differences in the performance of these methods reflect the failure of certain methods to account for intrinsic variation between biological replicates. This mechanistic understanding led to the discovery that the most frequently used methods can identify differentially expressed genes even in the absence of biological differences. These false discoveries are poised to mislead investigators. However, we show that false discoveries can be avoided using statistical methodologies that account for between-replicate variation. In summary, we expose the principles that underlie valid differential expression analysis in single-cell data, and provide a toolbox to implement relevant statistical methods for single-cell users.

## Results

### A ground-truth resource to benchmark single-cell differential expression

We aimed to compare available statistical methods for differential expression (DE) analysis based on their ability to generate biologically accurate results. We reasoned that performing this comparison in real datasets where the experimental ground truth is known would faithfully reflect differences in the performance of these methods, while avoiding the shortcomings of simulated data. We posited that the closest possible approximation to this ground truth could be obtained from matched bulk and scRNA-seq performed on the same population of purified cells, exposed to the same perturbations, and sequenced in the same laboratories. An extensive survey of the literature identified a total of eighteen ‘gold standard’ datasets that met these criteria (**Fig. 1a**)^13–16^.

**Fig. 1.**
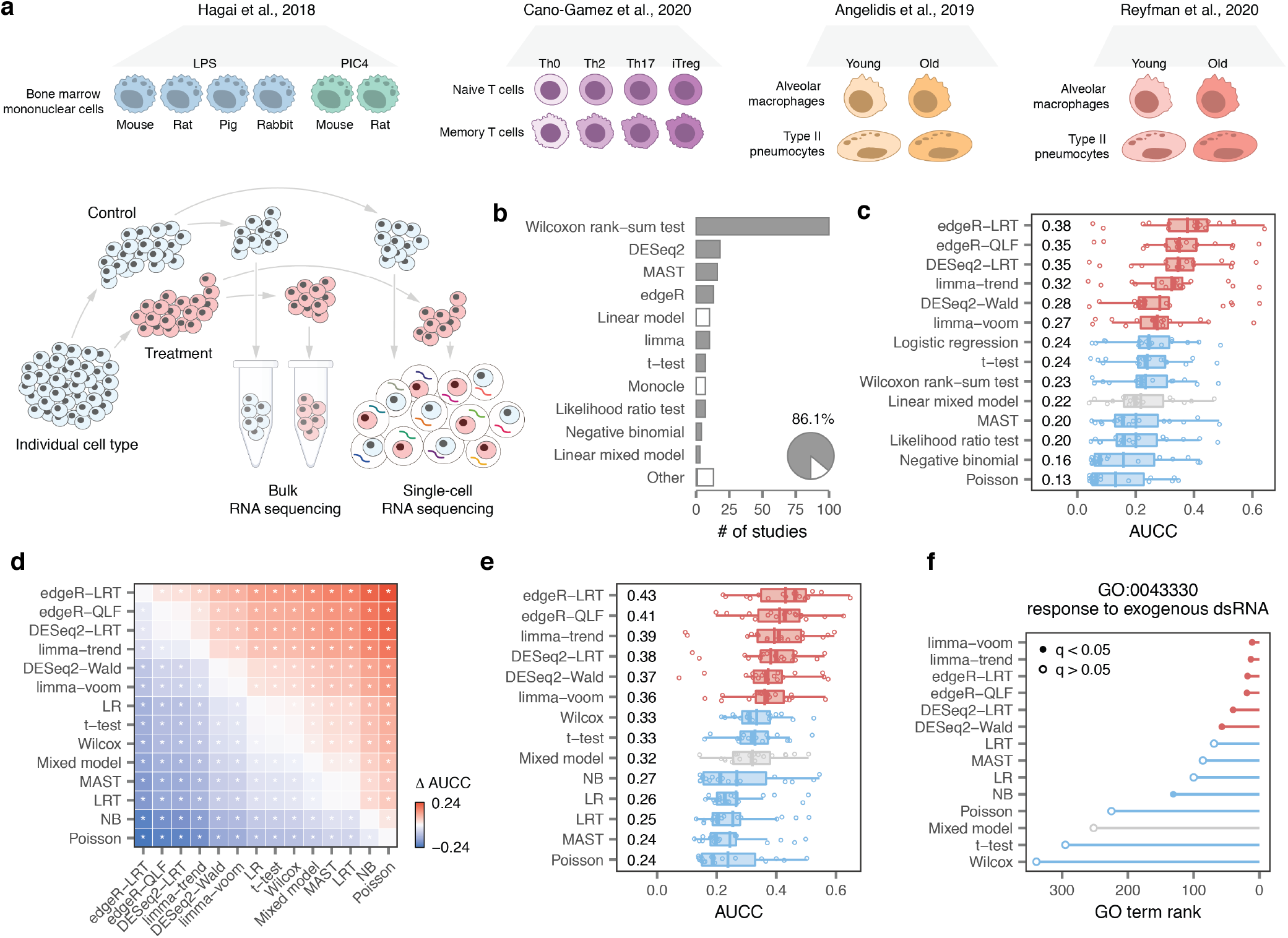
A systematic benchmark of differential expression in single-cell transcriptomics. **a,** Schematic overview of the eighteen ground-truth datasets analyzed in this study. **b,** Statistical methods for DE analysis employed in 500 recent scRNA-seq papers. Grey bars represent DE analysis methods included in this study. “Other” includes methods used in two or fewer studies. Inset pie chart shows the total proportion of recent scRNA-seq papers that employed DE analysis methods included in this study. **c,** Area under the concordance curve (AUCC) for fourteen DE methods in the eighteen ground-truth datasets shown in **a**. **d,** Mean difference in the AUCC (ΔAUCC) between the fourteen DE methods shown in c. Asterisks indicate comparisons with a two-tailed t-test p-value less than 0.05. **e,** AUCC of GO term enrichment, as evaluated using gene set enrichment analysis12, in the eighteen ground-truth datasets shown in **a**. **f,** Rank and statistical significance of the GO term GO:0043330 (“response to exogenous dsRNA”) in GSEA analyses of mouse bone marrow mononuclear cells stimulated with poly-I:C, a type of synthetic dsRNA, for four hours, using the output of fourteen DE methods.

This compendium allowed us to carry out a large-scale comparison of DE methods in experimental settings where the ground truth is known.

### Pseudobulk methods outperform generic and specialized single-cell DE methods

We selected a total of fourteen DE methods, representing the most widely used statistical approaches for single-cell transcriptomics, to compare (Methods). Together, these methods have figured in nearly 90% of recent studies (**Fig. 1b**). We evaluated the relative performance of each method based on the concordance between DE results in bulk versus scRNA-seq datasets. To quantify this concordance, we calculated the area under the concordance curve (AUCC) between the results of bulk versus scRNA-seq datasets^17,18^.

We compared the performance of the fourteen methods across the entire compendium of the eighteen gold standard datasets. This analysis immediately revealed that all six of the top-performing methods shared a common analytical property. These methods aggregated cells within a biological replicate, to form so-called ‘pseudobulks’, before applying a statistical test (**Fig. 1c**)^19^. In comparison, methods that compared individual cells performed poorly. The differences between pseudobulk and single-cell methods were highly significant (**Fig. 1d**), and robust to the methodology used to quantify concordance (**Supplementary Fig. 1a-d**). Moreover, comparisons to matching proteomics data^14^ revealed that pseudobulk methods also more accurately predicted changes in protein abundance (**Supplementary Fig. 1e-f**).

We asked whether the differences between DE methods could also impact the functional interpretation of transcriptomic experiments. For this purpose, we compared Gene Ontology (GO) term enrichment analyses in bulk versus scRNA-seq DE. We found that pseudobulk methods again more faithfully reflected the ground truth, as captured in the bulk RNA-seq (**Fig. 1e** and **Supplementary Fig. 1g**). For example, single-cell methods failed to identify the relevant GO term when comparing mouse phagocytes stimulated with poly(I:C)^13^, a synthetic double-stranded RNA (**Fig. 1f**).

### Single-cell DE methods are biased towards highly expressed genes

The unexpected superiority of pseudobulk methods compelled us to study the mechanisms that are responsible for their ability to recapitulate biological ground truth. To investigate these mechanisms, we formulated and tested several hypotheses that could potentially explain these differences in performance.

Previous studies demonstrated that inferences about DE are generally more accurate for highly expressed genes^20,21^. Measurements of gene expression in single cells are inherently sparse. By aggregating cells within each replicate, pseudobulk methods dramatically reduce the number of zeros in the data, especially for lowly expressed genes (**Fig. 2a**). Consequently, we initially hypothesized that the difference in accuracy between pseudobulk and single-cell methods could be explained by superior performance of pseudobulk methods among lowly expressed genes.

**Fig. 2.**
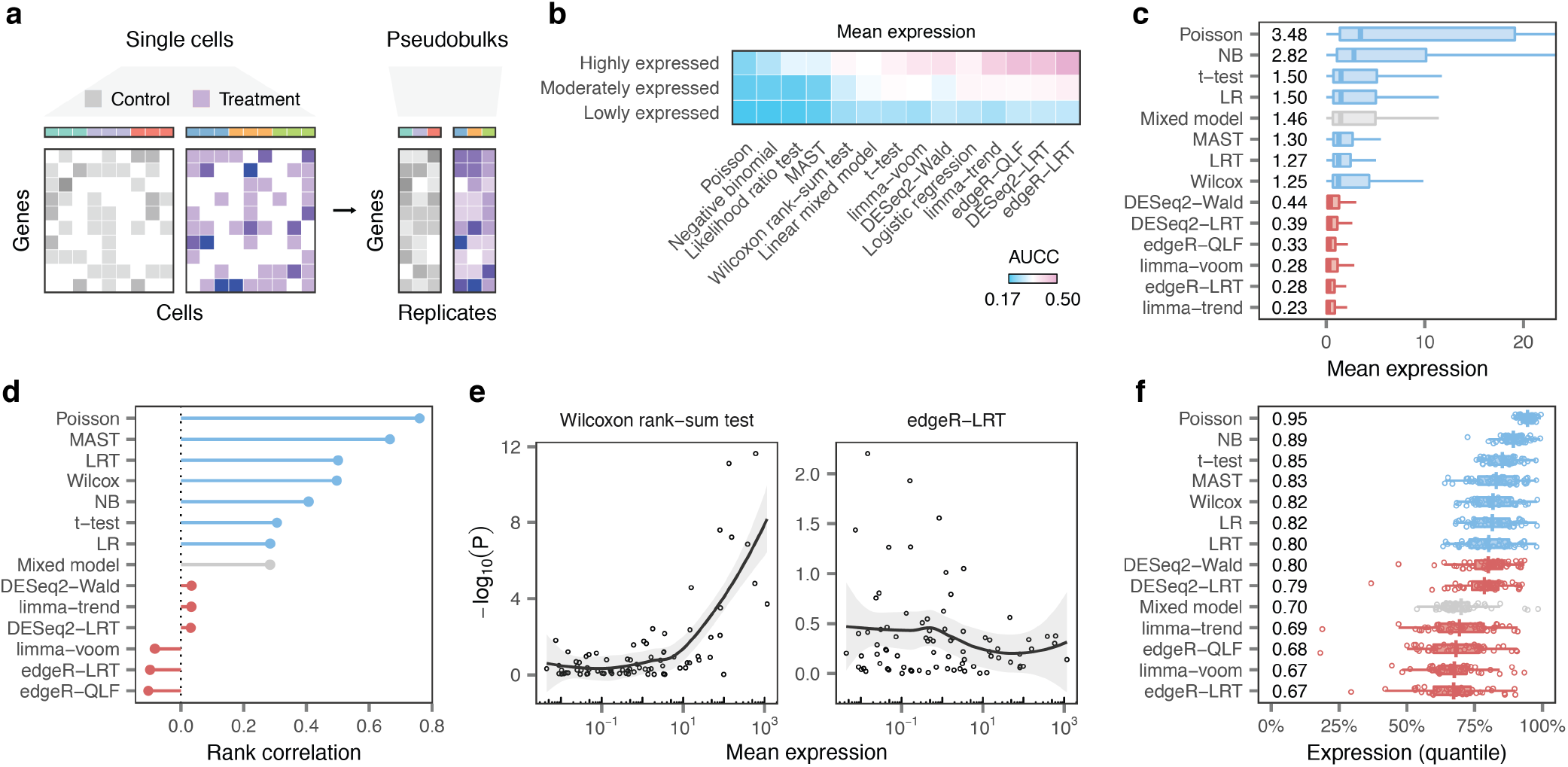
Single-cell DE methods are biased towards highly expressed genes. **a,** Schematic illustration of the creation of ‘pseudobulks’ from single-cell data by aggregating cells of a given type within each biological replicate. **b,** Mean AUCCs across eighteen ground-truth datasets after dividing the transcriptome into terciles of lowly, moderately, or highly expressed genes. **c,** Mean expression levels of the 100 top-ranked false-positive genes from each DE method. **d,** Spearman correlation between the mean expression of 80 ERCC spike-ins^13^ expressed in at least three cells and the −log_10_ p-value of differential expression assigned by each DE method. **e,** Scatterplots of mean ERCC expression vs. −log_10_ p-value for exemplary single-cell and pseudobulk DE methods. **f,** Mean expression levels of the 200 top-ranked genes from each DE method in a collection of 46 scRNA-seq datasets.

To test this hypothesis, we allocated genes into three equally sized bins, comprising lowly, moderately, and highly expressed genes. We then re-calculated the concordance between bulk and scRNA-seq DE within each bin. Contrary to our prediction, we observed minimal differences between pseudobulk and single-cell methods for lowly expressed genes (**Fig. 2b** and **Supplementary Fig. 2a**). Instead, the most pronounced differences between pseudobulk and single-cell methods emerged among highly expressed genes.

This unexpected result led us to ask whether single-cell DE methods produce systematic errors for highly expressed genes. To explore this possibility, we scrutinized the bulk datasets to identify genes falsely called as DE by each method within scRNA-seq data. We found that false positives identified by single-cell DE methods were more highly expressed than those identified by pseudobulk methods (**Fig. 2c** and **Supplementary Fig. 2b**). Conversely, false-negatives over-looked by single-cell DE methods tended to be lowly expressed (**Supplementary Fig. 2c-d**). Together, these findings implied a systematic tendency for single-cell methods to identify highly expressed genes as DE, even when their expression remained unchanged.

To validate this conclusion experimentally, we analyzed a dataset in which a population of synthetic mRNAs were spiked into each well containing a single cell^13,22^. Each of these single-cell libraries therefore contained equal concentrations of each synthetic mRNA. We found that single-cell methods incorrectly identified many abundant spike-ins as DE (**Fig. 2d-e** and **Supplementary Fig. 2e-f**). In contrast, pseudobulk methods avoided this bias.

We then asked whether this bias was universal in single-cell transcriptomics. We assembled a compendium of 46 scRNA-seq datasets that encompassed disparate species, cell types, technologies, and biological perturbations (**Supplementary Fig. 3**). We found that single-cell DE methods displayed a systematic preference for highly expressed genes across the entire compendium (**Fig. 2f**).

Together, these experiments suggest that the inferior performance of single-cell methods can be attributed to their bias towards highly expressed genes.

### DE analysis of single-cell data must account for biological replicates

These findings implied that pseudobulk methods possess a common analytical property that allows them to avoid this bias. We conducted a series of experiments to identify this mechanism.

The statistical tools applied to identify DE genes in pseudobulk data (i.e., edgeR, DESeq2, and limma) have been refined over many years of development. We therefore asked whether these methods incorporate inherent advantages that are independent from the procedure of aggregating gene expression across cells. To test this possibility, we disabled the aggregation procedure and applied the pseudobulk methods to individual cells (**Fig. 3a**). Strikingly, this procedure abolished the superiority of the pseudobulk methods (**Fig. 3b** and **Supplementary Fig. 4a**). The emergence of a bias towards highly expressed genes paralleled this decrease in performance (**Fig. 3b** and **Supplementary Fig. 4b-c**).

**Fig. 3.**
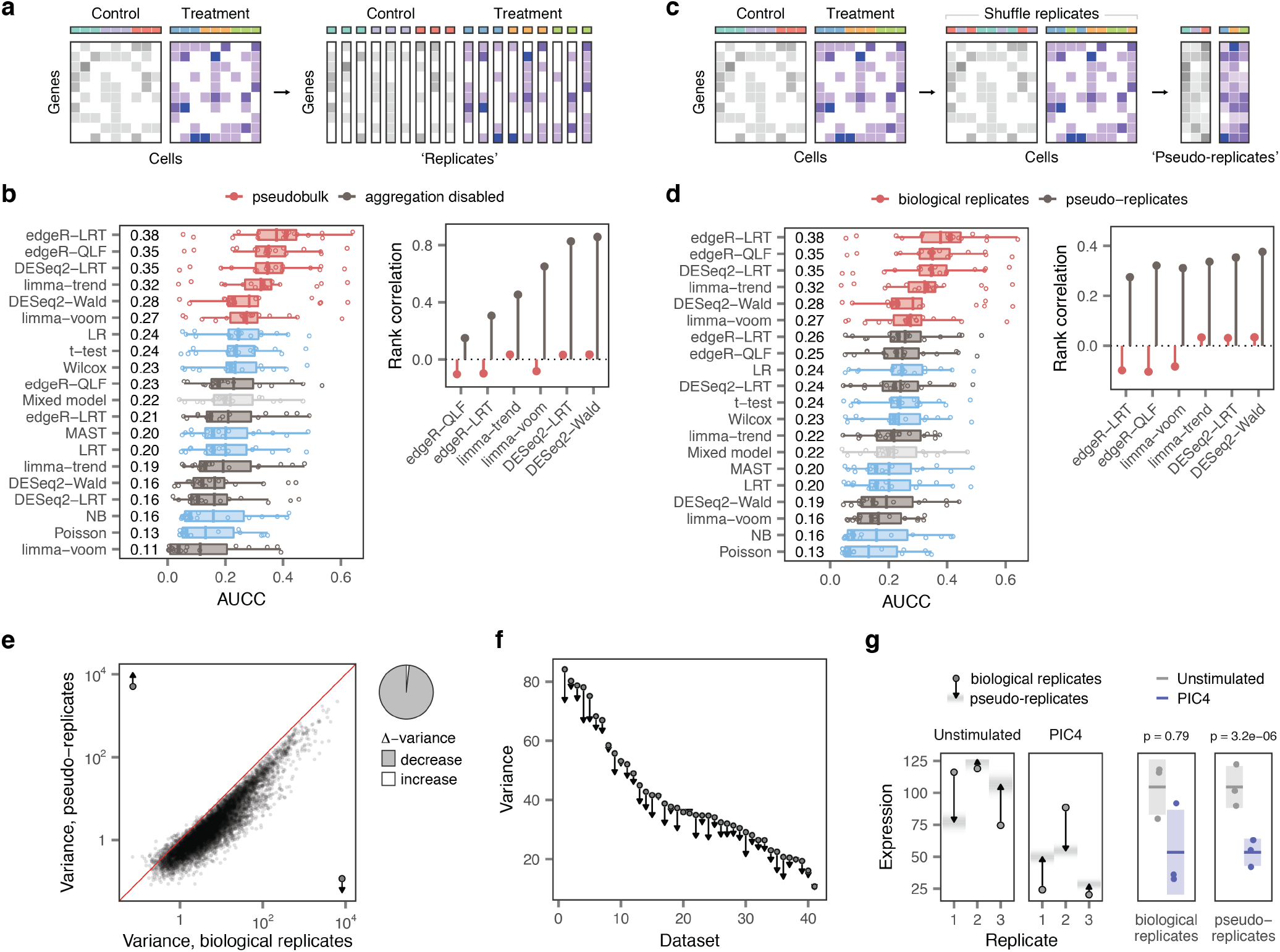
DE analysis of single-cell data must account for biological replicates. **a,** Schematic illustration of the experiment shown in **b**, in which the aggregation procedure was disabled and pseudobulk DE methods were applied to individual cells. **b,** Left, AUCC of the original fourteen DE methods, plus six pseudobulk methods applied to individual cells, in the eighteen ground-truth datasets. Right, Spearman correlation between ERCC mean expression^13^ and −log_10_ p-value assigned by six pseudobulk DE methods, before and after disabling the aggregation procedure. **c,** Schematic illustration of the experiment shown in **d**, in which the replicate associated with each cell was shuffled to produce ‘pseudo-replicates.’ **d,** Left, AUCC of the original fourteen DE methods, plus six pseudobulk methods applied to pseudo-replicates, in the eighteen ground-truth datasets. Right, Spearman correlation between ERCC mean expression^13^ and −log_10_ p-value assigned by six pseudobulk DE methods, before and after shuffling replicates to produce pseudo-replicates. **e,** Variance of gene expression in pseudobulks formed from biological replicates and pseudo-replicates in mouse bone marrow mononuclear cells stimulated with poly-I:C^13^. Shuffling the replicate associated with each cell produced a systematic decrease in the variance of gene expression. Right, pie chart shows the proportion of genes with increased or decreased variance in pseudo-replicates, as compared to biological replicates. **f,** Decreases in the variance of gene expression in pseudo-replicates as compared to biological replicates across 46 scRNA-seq datasets. Points show the mean variance in biological replicates; arrowheads show the mean variance in pseudo-replicates. **g,** Left, expression of the gene *Txnrd3* in biological replicates (points) and pseudo-replicates (arrowheads) from unstimulated cells and cells stimulated with poly-I:C, with the range of possible pseudo-replicate expression values shown as a density. Right, mean and variance of *Txnrd3* expression in biological replicates (left) and pseudo-replicates (right).

This result raised the possibility that the aggregation procedure itself was directly responsible for the superiority of pseudobulk methods. To evaluate this notion, we applied the aggregation procedure to random groups of cells, which produced a pseudobulk matrix composed of ‘pseudo-replicates’ (**Fig. 3c**). This experiment induced a similar decrease in the performance of pseudobulk methods, combined with the reemergence of a bias towards highly expressed genes (**Fig. 3d** and **Supplementary Fig. 3d-f**).

We sought to understand the common factors that could explain the decreased performance of pseudobulk methods in these two experiments. We recognized that both experiments entailed a loss of information about biological replicates. Aggregating random groups of cells to form pseudoreplicates, or ignoring replicates altogether in comparisons of single cells, both introduced a bias towards highly expressed genes and a corresponding loss of performance.

Within the same experimental condition, replicates exhibit inherent differences in gene expression, which reflect both biological and technical factors^23^. We reasoned that failing to account for these differences could lead methods to misattribute the inherent variability between replicates to the effect of the perturbation. To study this potential mechanism, we compared the variance in the expression of each gene in pseudobulks and pseudo-replicates. Initially, we performed this comparison in a dataset of bone marrow mononuclear cells stimulated with poly-I:C^13^. We found that shuffling the replicates produced a systematic decrease in the variance of gene expression, affecting 98.2% of genes (**Fig. 3e**). We next tested whether this decrease in variance occurred systematically across our compendium of 46 datasets. Every comparison displayed the same decrease in the variance of gene expression (**Fig. 3f**).

The decrease in the variance of gene expression led statistical tests to attribute small changes in gene expression to the effect of the perturbation. For instance, in the poly-I:C dataset, failing to account for the variable expression of *Txnrd3* across replicates led to the spurious identification of this gene as differentially expressed (**Fig. 3g**). Moreover, we found that highly expressed genes exhibited the largest decrease in variance in pseudo-replicates, thus explaining the bias of single-cell methods towards highly expressed genes (**Supplementary Fig. 4g-k**).

Together, this series of experiments exposed the principle underlying the unexpected superiority of pseudobulk methods. Statistical methods for differential expression must account for the intrinsic variability of biological replicates to generate biologically accurate results in single-cell data. Accounting for this variability allows pseudobulk methods to correctly identify changes in gene expression caused by a biological perturbation. In contrast, failing to account for biological replicates causes single-cell methods to systematically underestimate the variance of gene expression. This underestimation of the variance biases single-cell methods towards highly expressed genes, compromising their ability to generate biologically accurate results.

### False discoveries in single-cell DE

We realized that if failing to account for the variation between biological replicates could produce false discoveries in the presence of a real biological perturbation, then false discoveries might also arise in the absence of any biological difference. To test this possibility, we simulated single-cell data with varying degrees of heterogeneity between replicates (**Fig. 4a**). We randomly assigned each replicate to an artificial ‘control’ or ‘treatment’ group, and tested for DE between the two conditions. Strikingly, single-cell methods identified hundreds of DE genes in the absence of any perturbation (**Fig. 4b** and **Supplementary Fig. 5a**). Moreover, in line with our understanding of the mechanisms underlying the failure of single-cell DE methods, the genes that were falsely called as DE were those whose expression was most variable between replicates (**Fig. 4c** and **Supplementary Fig. 5b**). Pseudobulk methods abolished the false detection of DE genes. However, creating pseudo-replicates led to the reappearance of spurious DE genes (**Fig. 4b-c** and **Supplementary Fig. 5a-b**), further corroborating the requirements for accurate DE analyses. The number of false discoveries was reduced when additional replicates were introduced to the dataset (**Supplementary Fig. 5c**). In contrast, introducing additional cells to the simulated data only exacerbated the underlying problem (**Supplementary Fig. 5d**).

**Fig. 4.**
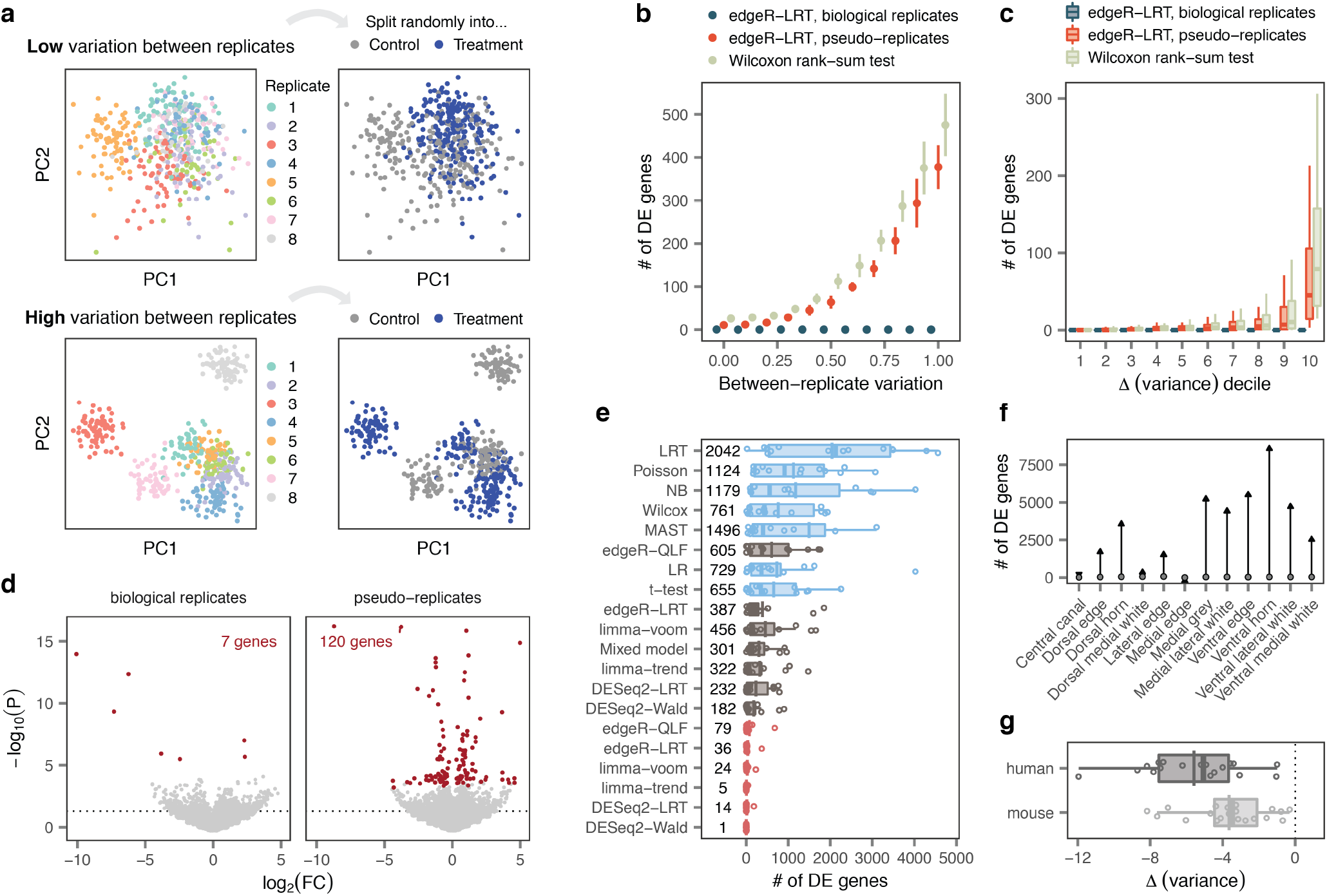
False discoveries in single-cell DE. **a,** Schematic illustration of simulation experiments. Single-cell RNA-seq datasets were simulated with varying degrees of heterogeneity between replicates. Replicates were then randomly assigned to either a ‘treatment’ or ‘control’ group, and DE analysis was performed between groups. **b,** Number of DE genes detected in stimulation experiments with varying degrees of heterogeneity between replicates by a representative single-cell DE method, a representative pseudobulk method, and the same pseudobulk method applied to pseudo-replicates. **c,** Number of DE genes detected by the tests shown in **b** for genes divided into deciles by the magnitude of the change in variance between biological replicates and pseudo-replicates (Δ-variance). **d,** Volcano plots showing DE between T cells from random groups of unstimulated controls drawn from Kang et al.^5^ using a representative pseudobulk method, edgeR-LRT, applied to biological replicates or pseudo-replicates. Discarding information about biological replicates leads to the appearance of false discoveries. **e,** Number of DE genes detected in comparisons of random groups of unstimulated controls from fourteen scRNA-seq studies with at least six control samples. **f,** Number of DE genes detected within spinal cord regions from control mice profiled by spatial transcriptomics^24^ using a representative pseudobulk method, edgeR-LRT (points), or a representative single-cell method, the Wilcoxon rank-sum test (arrowheads). **g,** Mean change in variance between biological replicates and pseudo-replicates for eighteen human and 20 mouse scRNA-seq datasets.

These findings compelled us to investigate whether similar false discoveries could arise in real single-cell data. To explore this possibility, we initially analyzed a dataset of human peripheral blood mononuclear cells (PBMCs) exposed to interferon^5^. We extracted the control samples that had not been exposed to interferon, and split them randomly into two groups. We then performed DE analysis. Failing to account for the intrinsic variability of biological replicates produced hundreds of DE genes between randomly assigned replicates (**Fig. 4d** and **Supplementary Fig. 6a-b**).

Unsettled by this appearance of false discoveries, we asked whether this observation reflected a universal pitfall. To address this concern comprehensively, we identified a total of fourteen datasets that included at least six replicates in the control condition. As in the previous experiment, we split these unperturbed samples randomly into synthetic control and treatment groups, before conducting DE analyses between these two groups. This systematic analysis confirmed that single-cell methods produced a systematic excess of false positives compared to pseudobulk methods (**Fig. 4e**). The resulting DE genes were enriched for hundreds of Gene Ontology (GO) terms, despite a complete absence of biological perturbation (**Supplementary Fig. 6c**). Moreover, we again confirmed that the genes falsely identified as DE corresponded to those with the highest variability between replicates (**Supplementary Fig. 6d**).

Together, these experiments exposed a fundamental pitfall for DE analysis in single-cell transcriptomics. We intuited, however, that this pitfall could afflict any technology in which many observations are obtained from each biological replicate. For example, we anticipated that false discoveries would also emerge in spatial transcriptomics data^25^. To test this prediction, we analyzed a spatial transcriptomics dataset that profiled spinal cords from a model of amyotrophic lateral sclerosis (ALS)^24^. We randomly partitioned data from control mice into two groups, and performed DE within each region of the spinal cord. Statistical methods that failed to account for variability between biological replicates identified thousands of DE genes within each region (**Fig. 4f** and **Supplementary Fig. 6e**). In contrast, pseudobulk methods abolished these false discoveries.

These experiments demonstrated that the variability between biological replicates can confound the identification of genes affected by a biological perturbation. Many of the factors that produce this variability between replicates can be minimized in animal models, including the genetic background, environment, intensity and timing of the biological perturbation, and sample processing. In contrast, these sources of variation are inherently more difficult to control in experiments involving human subjects. This distinction raised the possibility that single-cell studies of human tissue would exhibit greater variability between biological replicates, and consequently, would be more vulnerable to false discoveries. To evaluate this possibility systematically, we calculated the variability between replicates within 41 human and mouse scRNA-seq datasets. In agreement with our hypothesis, we detected significantly more variability between replicates in the human datasets (**Fig. 4g**). While we show that accounting for biological replicates is critical for any DE analysis, this result stresses the paramount importance of addressing this issue in single-cell studies of human tissue.

## Discussion

Accurate DE analysis in single-cell transcriptomics is required to dissect the transcriptional programs underlying the multifaceted responses to disease, trauma, and experimental manipulations. Despite the importance of statistical methods for DE analyses, the principles that determine their performance have remained elusive. Here, we demonstrate that the central principle underlying valid DE analysis is the ability of statistical methods to account for the intrinsic variability of biological replicates. Accounting for this variability dictates the biological accuracy of statistical methods. Conversely, methods that fail to account for the variability of biological replicates can produce hundreds of false discoveries in the absence of any biological difference.

Investigators study single cells to understand more general principles underlying the response to a biological perturbation. Clarifying these principles requires statistical inferences that generalize beyond the individual cells that constitute any particular dataset. Our results demonstrate that by performing a statistical inference at the level of individual cells, single-cell DE methods conflate variability between biological replicates with the effect of a biological perturbation. The presence of variability between replicates is an intrinsic biological phenomenon that cannot be avoided, and accordingly affects other high-dimensional assays, including spatial transcriptomics. Moreover, this variability is greatest in studies of human tissue, suggesting that inference at the level of biological replicates is critical to understand the cellular and molecular basis for human disease.

Our experiments established that accounting for variation between biological replicates dictated the performance of single-cell DE methods. We were therefore puzzled by the unsatisfying performance of a linear mixed model. By explicitly modeling variation both within and between biological replicates, mixed models should benefit from increased statistical power compared to pseudobulk methods^10^. To clarify this discrepancy, we evaluated eight additional Poisson or negative binomial generalized linear mixed models (GLMMs; **Supplementary Fig. 7a-b**). In datasets of 25-50 cells, GLMMs could produce accurate results under very specific parameter combinations. However, in datasets comprising 500 or more cells, their performance converged to that of pseudobulk DE methods. Moreover, the computational resources required to fit the best-performing GLMMs were enormous. Even in downsampled datasets, DE analysis of a single cell type took an average of 13.5 hours (**Supplementary Fig. 7c-d**). In contrast, pseudobulk methods required only minutes per cell type in our compendium of 46 datasets (**Supplementary Fig. 7e-f**). These observations suggest that, in practice, pseudobulk approaches provide an excellent trade-off between speed and accuracy for singlecell DE analysis.

Our results demonstrate that single-cell DE methods are poised to produce false discoveries. This understanding contrasts with the current default methods in the most widely used analysis packages in the field^26,27^. Our results stress the urgent need for a paradigm shift in the statistical methods that are used for DE analyses. To catalyze this transition, we implement all of the methods tested here in an R package that can be run on a laptop computer (**Supplementary Software 1**).

## Methods

### Literature review

To identify which statistical methods for DE analysis have been most commonly used within the field, we conducted an extensive literature review. We annotated the statistical method used to perform DE analysis across experimental conditions within cell types for each publication included in a large database of scRNA-seq studies^28^. Because this database spans a large period of time, and we wanted to establish which methods for DE analysis are in current use, we limited our analysis to the 500 most recently published studies. We did not annotate methods used to identify genes differentially expressed between cell types (i.e., marker gene identification), as this problem presents a distinct set of statistical challenges^9,29^.

### Ground-truth datasets

Previous benchmarks of DE analysis methods for single-cell transcriptomics have relied heavily on simulated data, or else have compared the results of different methods in scenarios where no ground truth was available^9,18^. We reasoned that the best possible approximation to the biological ground truth in a scRNA-seq experiment would consist of a matched bulk RNA-seq dataset in the same purified cell type, exposed to the same perturbation under identical experimental conditions, and sequenced in the same laboratory. We surveyed the literature to identify such matching single-cell and bulk RNA-seq datasets, which led us to compile a resource of eighteen ground truth datasets from four publications^13–16^. Datasets of mouse, rat, pig, and rabbit bone marrow-derived mononuclear phagocytes stimulated with either lipopolysaccharide or poly-I:C for 4 h were obtained from Hagai et al.^13^ Datasets of naive or memory T cells stimulated for 5 d with anti-CD3/anti-CD28 coated beads in the presence or absence of various combinations of cytokines (Th0: anti-CD3/anti-CD28 alone; Th2: IL-4, anti-IFN*γ*; Th17: TGR*β*, IL6, IL23, IL1*β*, anti-IFN*γ*, anti-IL4; iTreg: TGF,*β*, IL2) were obtained from Cano-Gamez et al.^14^ We additionally obtained label-free quantitative proteomics data for the same comparisons from this study. Datasets of alveolar macrophages and type II pneumocytes from young (3 m) and old (24 m) mice were obtained from Angelidis et al.^15^ Datasets of alveolar macrophages and type II pneumocytes from patients with pulmonary fibrosis and control individuals were obtained from Reyfman et al.^16^

### Differential expression analysis methods

We compared fourteen statistical methods for DE analysis of single-cell transcriptomics data on their ability to recover ground-truth patterns of DE, as established through bulk RNA-seq analysis of matching cell populations. These fourteen methods comprised seven statistical tests that compared gene expression in individual cells (“singlecell methods”); six tests that aggregated cells within a biological replicate to form pseudobulks before performing statistical analysis (“pseudobulk methods”); and a linear mixed model.

The seven single-cell methods analyzed here included a t-test, a Wilcoxon rank-sum test, logistic regression^30^, negative binomial and Poisson generalized linear models, a likelihood ratio test^31^, and the two-part hurdle model implemented by MAST^8^. The implementation provided in the Seurat function ‘FindMarkers’ was used for all seven tests, with all filters (‘min.pct’, ‘min.cells.feature’, and ‘logfc.threshold’) disabled. In addition, we implemented a linear mixed model within Seurat, using the ‘lmerTest’ R package to optimize the restricted maximum likelihood and obtain p-values from the Satterthwaite approximation for degrees of freedom. We observed that some statistical tests returned a large number of p-values below the double precision limit in R (approximately 2 × 10^−308^), potentially confounding the calculation of the concordance metrics described below. To avoid this pitfall, we modified the Seurat implementation to also return the value of the test statistic from which the p-value was derived. The modified version of Seurat 3.1.5 used to perform all single-cell DE analyses reported in this study is available from http://github.com/jordansquair/Seurat.

The pseudobulk methods employed the DESeq2^32^, edgeR^33^, and limma^34^ packages for analysis of aggregated read counts. Briefly, for cells of a given type, we first aggregated reads across biological replicates, transforming a genes-by-cells matrix to a genes-by-replicates matrix using matrix multiplication. For DESeq2, we used both a Wald test of the negative binomial model coefficients (DESeq2-Wald) as well as a likelihood ratio test compared to a reduced model (DESeq2-LRT) to compute the statistical significance. For edgeR, we used both the likelihood ratio test (edgeR-LRT)^35^ as well as the quasi-likelihood F-test approach (edgeR-QLF)^36^. For limma, we compared two modes: limma-trend, which incorporates the mean-variance trend into the empirical Bayes procedure at the gene level, and voom (limma-voom), which incorporates the mean-variance trend by assigning a weight to each individual observation^37^. Log-transformed counts per million values computed by edgeR were provided as input to limma-trend.

DE analysis of bulk RNA-seq datasets was performed with six methods (DESeq2-LRT, DESeq2-Wald, edgeR-LRT, edgeR-QLF, limma-trend, and limma-voom), except for the two pulmonary fibrosis datasets^16^; for these datasets, the raw bulk RNA-seq data from sorted cells could not be obtained, so only the results of the bulk DE analysis performed by the authors of the original publication were used. The AUCC and rank correlation were calculated for each bulk DE analysis method separately, and subsequently averaged over all six methods. DE analysis of normalized bulk proteomics data was performed using the moderated t-test implemented within limma, as in the original publication.

### Measuring concordance between single-cell and bulk RNA-seq

To evaluate the concordance between DE analyses of matched single-cell and bulk RNA-seq data, we computed two metrics, designed to evaluate the concordance between only the most highly ranked subset of DE genes and across the entire transcriptome, respectively. To calculate the first of these metrics, the area under the concordance curve (AUCC)^17,18^, we ranked genes in both the single-cell and bulk datasets in descending order by the statistical significance of their differential expression. Then, we created lists of the top-ranked genes in each dataset of matching size, up to some maximum size *k*. For each of these lists (that is, for the top-1 genes, top-2 genes, top-3 genes, and so on), we computed the size of the intersection between the single-cell and bulk DE genes. This procedure yielded a curve relating the number of shared genes between datasets to the number of top-ranked genes considered. The area under this curve was computed by summing the size of all intersections, and normalized to the range [0, 1] by dividing it by its maximum possible value, *k* × (*k* + 1) / 2. To evaluate the concordance of DE analysis, we used *k* = 500 except where otherwise noted, but found our results were insensitive to the precise value of *k*. To compute the second metric, the transcriptome-wide rank correlation, we multiplied the absolute value of the test statistic for each gene by the sign of its log-fold change between conditions, and then computed the Spearman correlation over genes between the singlecell and bulk datasets.

In addition to evaluating the consistency of DE analyses at the gene level, we also asked whether each DE method yielded broader patterns of functional enrichment that were similar between the single-cell and bulk datasets, allowing for some divergence in the rankings of individual genes. To address this question, we performed gene set enrichment analysis^12^ using the ‘fgsea’ R package^38^. GO term annotations for human and mouse (2019-12-09 release) were obtained from the Gene Ontology Consortium website. GO terms annotated to less than 10 genes or more than 1,000 genes within each dataset were excluded in order to mitigate the influence of very specific or very broad terms. Genes were ranked in descending order by the absolute value of the test statistic, and 10^6^ permutations were performed. To evaluate the concordance of GO term enrichment, we used *k* = 100, on the basis that fewer top-ranked GO terms are generally of interest than are top-ranked genes.

### Impact of mean expression

We initially hypothesized that differences between single-cell DE analysis methods could be attributed to their differing sensitivities towards lowly expressed genes. To explore this hypothesis, we performed the following analyses. First, we divided genes from the eighteen gold standard datasets into three equally sized bins on the basis of their mean expression, then re-calculated the AUCC as described above within each bin separately. Second, we inspected the properties of genes falsely called as DE in the single-cell data (false positives) or incorrectly inferred to be unchanging in the single-cell data (false negatives). To identify false positive genes, we used the bulk DE analysis to exclude genes called as DE at a false discovery rate of 10% from the matched single-cell results, then retained the 100 top-ranked remaining genes in the single-cell data. To identify false negative genes, we used the bulk DE analysis to identify genes called as DE at a false discovery rate of 10%, but with a false discovery rate exceeding 10% in the matched single-cell results, again retaining the 100 top-ranked such genes. For each of these genes, we computed both the mean expression level and the proportion of zero gene expression measurements. Third, we analyzed a Smart-seq2 dataset of human dermal fibroblasts stimulated with interferon-*β*, in which a mixture of synthetic RNAs was spiked into each individual cell^13^. We performed DE analysis on the synthetic spike-ins, then calculated the Spearman correlation between the mean expression level of each spike-in and the statistical significance of differential expression, as assigned by each single-cell DE method. Fourth, we assembled a compendium of 46 published scRNA-seq datasets, and asked whether the genes called as DE by each method tended to be more or less highly expressed across the entire compendium. Complete details on the preprocessing of these 46 datasets are provided below. Because each of these datasets were sequenced to different depths, and captured different total numbers of genes (depending on both the sequencing depth and the biological system under study), mean expression values were not directly comparable across datasets. To enable such a comparison, we first calculated the mean expression for each gene, then converted this value into the quantile of mean expression using the empirical cumulative distribution function. We then calculated the mean expression quantile of the 200 top-ranked genes from each method in each of the 46 datasets.

### Dissecting pseudobulk DE methods

To understand the principles underlying the improved performance of the six pseudobulk DE methods, we performed the following analyses. First, we disabled the aggregation procedure that led to the creation of pseudobulks (that is, we treated each individual cell as its own replicate), then performed an identical DE analysis of individual cells. For each DE method, we then re-calculated both the AUCC and the bias towards highly expressed genes, as quantified by (i) the rank correlation to mean-spike in expression, and (ii) the expression quantile across 46 scRNA-seq datasets. Second, we aggregated random groups of cells into ‘pseudo-replicates’ by randomizing the replicate associated with each cell. We then again re-calculated both the AUCC and the bias towards highly expressed genes.

These experiments led us to suspect that discarding information about the inherent variability of biological replicates caused both the bias towards highly expressed genes and the attendant decrease in performance. To test this hypothesis, we compared the variance of gene expression in pseudobulks and pseudo-replicates. For each gene, we calculated the difference in variance (‘Δ-variance’) between pseudobulks and pseudo-replicates. We initially visualized the Δ-variance in an exemplary dataset, consisting of mouse bone marrow mononuclear cells stimulated with poly-I:C^13^. Subsequently, we calculated the mean Δ-variance across all genes in each of the 46 datasets in our scRNA-seq compendium, observing a decrease in the variance in all 46 cases. To clarify the relationship between the Δ-variance and mean gene expression, we computed the correlation between Δ-variance and mean expression, first in the poly-I:C dataset and then across all 46 datasets in the compendium. We observed a significant negative correlation, confirming that the variance of highly expressed genes is disproportionately underestimated when discarding information about biological replicates. We performed a similar analysis correlating the original variance of gene expression to the Δ-variance, demonstrating that the variance of the most variable genes is disproportionately underestimated when discarding information about biological replicates. However, in partial correlation analyses, only gene expression variance remained correlated with Δ-variance, implying that failing to account for biological replicates induces a bias towards highly expressed genes because these genes are also more variably expressed. **Supplementary Fig. 4h-i** employ the signed pseudologarithm transformation from the ‘ggallin’ R package to visualize the Δ-variance.

### Simulation studies

Our understanding of the importance of accounting for variability between biological replicates led us to ask whether failing to account for biological replication could lead to the appearance of false discoveries in the absence of a perturbation. To test this hypothesis, we simulated scRNA-seq data with no biological effect, in which we systematically varied the degree of heterogeneity between replicates. Simulations were performed using the ‘Splatter’ R package^39^, with simulation parameters estimated from the Kang et al. dataset^5^ using the ‘splatEstimate’ function. Populations of between 100 and 2,000 cells were simulated, with between 3 and 20 replicates per condition. DE of varying magnitudes was simulated between replicates by varying the location parameter of the DE factor log-normal distribution (‘de.facLoc’) between 0 and 1, treating each replicate as its own group, and the total proportion of DE genes (‘de.prob’) set to 0.5. Then, half of the replicates were randomly assigned to an artificial ‘treatment’ condition and the remaining half to a ‘control’ condition, and DE analysis was performed between the treatment and control groups. Except where otherwise noted, plots show results from a simulated population of 500 cells, with three replicates per condition.

### Analysis of published scRNA-seq control groups

To confirm that the trends observed in simulation studies were reflective of experimental datasets, we performed a similar analysis using published scRNA-seq data. Within our compendium, we identified a total of fourteen studies with control groups that included six or more samples^5,6,16,40–50^. Details on the preprocessing of each of these datasets are provided below. For each of these studies, we split the control group randomly into artificial ‘control’ and ‘treatment’ groups, and performed DE analysis. In addition to computing the total number of DE genes, we identified GO terms enriched among DE genes using a hypergeometric test. We also performed a similar analysis for one spatial transcriptomics dataset^24^, identifying DE genes between random groups of control mice with bar-codes grouped by spinal cord region rather than cell type. Spatial transcriptomics data was downloaded from the supporting website at https://als-st.nygenome.org. Only data from wild-type mice was retained for the analysis. Last, we hypothesized that scRNA-seq studies of human tissues would display more heterogeneity between replicates than studies of animal models, where factors such as genotype, environment, and perturbation can be precisely controlled. To test this hypothesis, we computed the mean Δ-variance across all genes in the 38 human or mouse scRNA-seq datasets in our compendium (*n* = 18 human datasets and 20 mouse datasets).

### Mixed models

Having established that the performance of DE methods is contingent on their ability to account for biological replicates, we asked why mixed models failed to match the performance of pseudobulk methods. In addition to the linear mixed model described above, we implemented generalized linear mixed models (GLMMs) based on the negative binomial or Poisson distributions, adapting implementations provided in the ‘muscat’ R package^9^. For each of these models, we evaluated the impact of incorporating the library size factors as an offset term, and compared the Wald test of model coefficients to a likelihood ratio test against a reduced model, yielding a total of four GLMMs from each distribution. The enormous computational requirements of the GLMMs prevented us from evaluating these models in the full ground truth datasets; instead, we analyzed a series of downsampled datasets, each containing between 25 and 1,000 cells. To quantify the computational resources required by each DE method, we monitored peak memory usage using the ‘peakRAM’ R package, and the base R function ‘system.time’ to record wall time.

### Preprocessing and analysis of published single-cell datasets

We assembled a compendium of 46 published singlecell or single-nucleus RNA-seq studies (**Supplementary Fig. 3**), and performed DE analyses across this compendium to establish the generality of our conclusions. For publications containing more than one comparison, only a single comparison was retained, as described in detail below. We retained the comparison involving the greatest number of cells, and used the most fine-grained cell type annotations provided by the authors of the original studies. When count matrices did not use gene symbols, the provided identifiers were mapped to gene symbols, and counts summed across genes mapping to identical symbols. Only cell types with at least three cells in each condition were subjected to DE analysis, and genes detected in less than three cells were removed.

*Angelidis et al., 2019*^15^. scRNA-seq data from young and aged mouse lung (3 m and 24 m, respectively), as well as matching bulk data from two purified cell types, was obtained from GEO (GSE124872). Metadata was obtained from GitHub (https://github.com/theislab/2018_Angelidis). Cells with missing cell type annotations were removed from the single-cell data. DE analysis was performed by comparing cells from young and old mice.

*Arneson et al., 2018*^51^. scRNA-seq data from the hippocampus of mice after a mild traumatic brain injury (mTBI), delivered using a mild fluid percussion injury model, and matched controls was obtained from GEO (accession: GSE101901). Metadata, including cell type annotations, were provided by the authors. DE analysis was performed by comparing cells from mTBI and control mice.

*Avey et al., 2018*^52^. scRNA-seq data from the nucleus accumbens of mice treated with morphine for 4 h and saline-treated controls was obtained from GEO (accession: GSE118918). Cells identified as doublets and non-unique barcodes were removed. Metadata, including cell type annotations, were provided by the authors. DE analysis was performed by comparing cells from morphine- and saline-treated mice.

*Aztekin et al., 2019*^53^. scRNA-seq data from regeneration-competent (NF stage 40-41) Xenopus laevis tadpoles was obtained from ArrayExpress (E-MTAB-7716). DE analysis was performed by comparing cells from tadpoles at 1 d post-amputation to control tadpoles.

*Bhattacherjee et al., 2019*^54^. scRNA-seq data from the prefrontal cortex of mice exposed to a cocaine withdrawal paradigm was obtained from GEO (accession: GSE124952). DE analysis was performed by comparing cells at the 15 d post-withdrawal timepoint from cocaine- or saline-treated mice.

*Brenner et al., 2020*^55^. snRNA-seq data from the prefrontal cortex of alcoholic and control individuals was obtained from GEO (accession: GSE141552). Metadata, including cell type annotations, were provided by the authors. DE analysis was performed by comparing nuclei from alcoholic and control individuals.

*Cano-Gamez et al., 2020*^14^. scRNA-seq data from naive and memory T cells, stimulated with anti-CD3/anti-CD28 coated beads in the presence or absence of various combinations of cytokines, was obtained from the supporting website (https://www.opentargets.org/projects/effectorness). Matching bulk RNA-seq and proteomics data was obtained from the same source. For the analyses presented as part of the compendium of 46 datasets, DE analysis was performed by comparing iTreg and control cells.

*Cheng et al., 2019*^56^. scRNA-seq data from intestinal crypt cells in wild-type and Hmgcs2 knockout mice was obtained directly from the authors of the original publication. DE analysis was performed by comparing wild type and KO mice.

*Co et al., 2020*^57^. scRNA-seq data of sorted cells from Drd1a-tdTomato+ control and Foxp2 KO mice was obtained from GEO (accession: GSE130653). Cell type annotations were provided by the authors. Cell types annotated as ‘Low quality’ were removed prior to further analysis. DE analysis was performed by comparing WT and Foxp2 KO mice.

*Crowell et al., 2020*^9^. snRNA-seq data from the prefrontal cortex of mice peripherally stimulated with lipopolysaccharide (LPS) and control mice was obtained from the Bioconductor package ‘muscData’, using the ‘Crowell19_4vs4’ function. DE analysis was performed by comparing nuclei from LPS-treated and control mice.

*Davie et al., 2018*^58^. scRNA-seq data from the brains of flies of varying ages, sexes, and genotypes was obtained from the supporting website (http://scope.aertslab.org, file ‘Aerts_Fly_AdultBrain_Filtered_57k.loom’). Cells marked as ‘Unannotated’ were removed. DE analysis was performed by comparing cells from DGRP-551 and W^1118^ flies.

*Denisenko et al., 2020*^59^. scRNA-seq data from human kidneys subjected to varying dissociation methods and cell fixation techniques was obtained from GEO (accession: GSE141115). Metadata, including cell type annotations, was obtained from the supporting information files accompanying the published manuscript. DE analysis was performed by comparing cells fixed with methanol and freshly dissociated cells, both at −20 °C.

*Der et al., 2019*^60^. scRNA-seq data of skin samples from patients with lupus nephritis (LN) and healthy controls was obtained from ImmGen (accession: SDY997; experiment: EXP15077). Cell type annotations were obtained from the authors of the original manuscript. Other metadata, including biological replicate and experimental condition annotations for each individual cell, was obtained from the supporting information files accompanying the published manuscript. DE analysis was performed by comparing cells from patients with LN and healthy controls.

*Goldfarbmuren et al., 2020*^48^. scRNA-seq data of tracheal epithelial cells from smokers and non-smokers was obtained from GEO (accession: GSE134174). Patients designated as ‘excluded’ were removed prior to downstream analysis. DE analysis was performed by comparing cells from smokers and non-smokers.

*Grubman et al., 2019*^43^. snRNA-seq data from the entorhinal cortex of patients with Alzheimer’s disease and matched controls was obtained from the supporting website (http://adsn.ddnetbio.com). Nuclei annotated as ‘undetermined’ or ‘doublet’ were removed. DE analysis was performed by comparing nuclei from patients with Alzheimer’s disease and controls.

*Gunner et al., 2019*^61^. scRNA-seq data from the mouse barrel cortex before or after whisker lesioning was obtained from GEO (accession: GSE129150). Cell types not included in Supplementary Fig. 10 of the original publication were removed. DE analysis was performed by comparing cells from lesioned and control mice.

*Haber et al., 2017*^62^. scRNA-seq data from epithelial cells of the mouse small intestine in healthy mice and after ten days of infection with the parasitic helminth Heligmosomoides polygyrus was obtained from GEO (accession: GSE92332), using the Drop-seq data collected by the original publication. DE analysis was performed by comparing cells from infected and uninfected mice.

*Hagai et al., 2018*^13^. scRNA-seq data of bone marrow-derived mononuclear phagocytes from four different species (mouse, rat, pig, and rabbit) exposed to lipopolysaccharide (LPS) or poly-I:C for two, four, or six hours was obtained from ArrayExpress (accession: E-MTAB-6754). Matching bulk RNA-seq data was also obtained from ArrayExpress (accession: E-MTAB-6773). Finally, scRNA-seq data from human dermal fibroblasts stimulated with interferon-*β* for two or six hours, in which the ERCC mixture of synthetic mRNAs was spiked in alongside every cell, was obtained from ArrayExpress (accession: E-MTAB-7051). Counts were summed across technical replicates of the same biological samples. For the analyses presented as part of the compendium of 46 datasets, DE analysis was performed by comparing rabbit cells stimulated with LPS for 2 h and control cells. DE analysis of the spike-in dataset was performed by comparing cells stimulated for 2 h and 6 h.

*Hashimoto et al., 2019*^63^. scRNA-seq data of peripheral blood mononuclear cells from human supercentenarians and younger controls was obtained from the supporting website (http://gerg.gsc.riken.jp/SC2018). Metadata, including cell type annotations, were provided by the authors. DE analysis was performed by comparing cells from supercentenarians and younger controls.

*Hrvatin et al., 2018*^41^. scRNA-seq data from the visual cortex of mice housed in darkness, then exposed to light for 0 h, 1 h, or 4 h was obtained from GEO (accession: GSE102827). Cell types labeled as ‘NA’ were removed from downstream analyses. DE analysis was performed by comparing cells from mice stimulated with light for 4 h to control mice.

*Hu et al., 2017*^64^. snRNA-seq data from the cerebral cortex of mice after pentylenetetrazole (PTZ)-induced seizure and saline-treated controls was obtained from the Google Drive folder accompanying the original publication (https://github.com/wulabupenn/Hu_MolCell_2017). DE analysis was performed by comparing cells from PTZ- and saline-treated mice.

*Huang et al., 2019*^44^. scRNA-seq data from the colon of pediatric patients with colitis and inflammatory bowel disease and matched controls was obtained from the supporting website (https://zhanglaboratory.com/research-data/). DE analysis was performed by comparing cells from patients with colitis and healthy controls.

*Jaitin et al., 2019*^65^. scRNA-seq data from white adipose tissue of mice fed either a high-fat diet or normal chow for six weeks were obtained from the Bitbucket repository accompanying the original publication (https://bitbucket.org/account/user/amitlab/projects/ATIC). Metadata, including cell type annotations, were provided by the authors. DE analysis was performed by comparing cells from high-fat diet and normal chow-fed mice.

*Jakel et al., 2019*^66^. snRNA-seq data of oligodendrocytes from patients with multiple sclerosis and matched controls was obtained from GEO (accession: GSE118257). DE analysis was performed by comparing nuclei from individuals with multiple sclerosis versus matched controls.

*Kang et al., 2018*^5^. scRNA-seq data from peripheral blood mononuclear cells (PBMCs) stimulated with recombinant IFN-*β* for 6 h and unstimulated PBMCs was obtained from GEO (accession: GSE96583). Doublets and unclassified cells were removed. DE analysis was performed by comparing IFN-stimulated and unstimulated cells.

*Kim et al., 2019*^67^. scRNA-seq data from the ventromedial hypothalamus of mice exposed to a range of behavioral stimuli and control mice was obtained from the Mendeley repository accompanying the original publication. Cell type annotations were provided directly by the authors. DE analysis was performed by comparing cells from animals engaging in aggressive behaviour to the common population of control animals.

*Kotliarov et al., 2020*^68^. scRNA-seq data of peripheral blood mononuclear cells from subjects who were subsequently given an influenza vaccination were obtained from Figshare (https://doi.org/10.35092/yhjc.c.4753772). DE analysis was performed by comparing cells from high and low responders to the influenza vaccination, as categorized by the authors.

*Madissoon et al., 2020*^69^. scRNA-seq data from esophagus, lung, and spleen samples after varying durations of cold storage was obtained from the study website (https://cellgeni.cog.sanger.ac.uk/tissue-stability/). DE analysis was performed by comparing cells from samples preserved for 12 h and fresh samples.

*Mathys et al., 2019*^6^. snRNA-seq data from the prefrontal cortex of patients with Alzheimer’s disease and matched controls was obtained from Synapse (accession: syn18681734). Patient data and additional metadata were also obtained from Synapse (accessions: syn3191087 and syn18642926, respectively). DE analysis was performed by comparing nuclei from patients with Alzheimer’s disease and controls.

*Nagy et al., 2020*^49^. snRNA-seq data from the dorsolateral prefrontal cortex of patients with major depressive disorder (MDD) and matched controls was obtained from GEO (accession: GSE144136). DE analysis was performed by comparing nuclei from patients with MDD and controls.

*Nault et al., 2021*^70^. snRNA-seq data from the livers of mice gavaged with 2,3,7,8-tetrachlorodibenzo-p-dioxin or sesame oil vehicle was obtained from GEO (accession: GSE148339). DE analysis was performed by comparing nuclei from treated and vehicle livers.

*Ordovas-Montanes et al., 2018*^71^. scRNA-seq data from ethmoid sinus cells of patients with chronic rhinosinusitis (CRS), with and without nasal polyps, from Supplementary Table 2 of the original publication. DE analysis was performed by comparing cells from patients with polyposis and non-polyposis CRS.

*Reyes et al., 2020*^72^. scRNA-seq data of peripheral blood mononuclear cells from patients with sepsis and healthy controls was obtained from the Broad Institute’s Single Cell Portal (accession: SCP548). DE analysis was performed by comparing cells from individuals with bacterial sepsis and controls.

*Reyfman et al., 2019*^16^. scRNA-seq data from the lungs of patients with pulmonary fibrosis and healthy controls was obtained from GEO (accession: GSE122960). Metadata, including cell type annotations, was provided by the authors. One sample (“Cry-obiopsy_01”) was removed as it was sequenced separately from the rest of the experiment. The results of DE analysis of bulk RNA-seq data, comparing purified AT2 cells or alveolar macrophages from patients with pulmonary fibrosis and healthy controls, were obtained from the supporting information accompanying the original publication. DE analysis was performed by comparing cells from patients with pulmonary fibrosis and controls.

*Rossi et al., 2019*^45^. scRNA-seq data from the hypothalamus of mice fed either a high-fat diet or normal chow for between 9-16 weeks was obtained directly from the authors, in the form of a processed Seurat object. Cells annotated as ‘unclassified’ were removed. DE analysis was performed by comparing cells from high-fat diet and normal chow-fed mice.

*Sathyamurthy et al., 2018*^42^. snRNA-seq data from the spinal cord parenchyma of adult mice exposed to formalin or matched controls was obtained from GEO (accession: GSE103892). Cell types with blank annotations, or annotated as ‘discarded’, were removed. DE analysis was performed by comparing cells from mice exposed to formalin and control animals.

*Schafflick et al., 2020*^73^. scRNA-seq data of peripheral blood mononuclear cells from individuals with multiple sclerosis and matched controls was obtained from GEO (accession: GSE138266). Metadata, including cell type annotations, was obtained from Github (https://github.com/chenlingantelope/MSscRNAseq2019). DE analysis was performed by comparing cells from individuals with multiple sclerosis and controls.

*Schirmer et al., 2019*^74^. snRNA-seq data from cortical and subcortical areas from the brains of patients with multiple sclerosis and control tissue from unaffected individuals was obtained from the web browser accompanying the original publication (https://cells.ucsc.edu/ms). DE analysis was performed by comparing cells from multiple sclerosis and control patients.

*Skinnider et al., 2021*^75^. snRNA-seq data from the spinal cords of mice with a spinal cord injury, some of which were exposed to epidural electrical stimulation to restore locomotion after paralysis, was obtained from GEO (accession: GSE142245). DE analysis was performed by comparing nuclei from paralyzed and walking mice.

*Tran et al., 2019*^47^. scRNA-seq data from the retinal ganglion of mice at various timepoints after an optic nerve crush injury, as well as uninjured controls, was obtained from GEO (accession: GSE137398). Metadata, including cell type annotations, was obtained from the Broad Institute’s Single-Cell Portal (accession: SCP509). DE analysis was performed by comparing cells from mice at 12 h post-injury and uninjured mice.

*Wagner et al., 2018*^76^. scRNA-seq data from zebrafish embryos between 14-16 hours post-fertilization, with either the chordin locus or a control locus (tyrosinase) disrupted by CRISPR-Cas9 knockout, was obtained from GEO (accession: GSE112294). DE analysis was performed by comparing cells from chordin- or tyrosinase-targeted embryos.

*Wang et al., 2020*^77^. scRNA-seq data from the ovaries of young and old cynomolgus monkeys was obtained from GEO (accession: GSE130664). Metadata, including cell type annotations, was obtained from the supporting information accompanying the original publication. Spike-ins were removed. DE analysis was performed by comparing cells from young and old primates.

*Wilk et al., 2020*^50^. scRNA-seq data of peripheral blood mononuclear cells from patients with COVID-19 and healthy controls was obtained from the supporting website (https://www.covid19cellatlas.org/). DE analysis was performed by comparing patients with COVID-19 and controls.

*Wirka et al., 2019*^78^. scRNA-seq data from the aortic root of mice fed a high-fat diet or normal chow for eight weeks from GEO (accession: GSE131776). Metadata, including cell type annotations, was provided by the authors, and unannotated cells were removed. DE analysis was performed by comparing cells from high-fat diet and normal chow-fed mice.

*Wu et al., 2017*^40^. scRNA-seq data from the amygdala of mice subjected to 45 min of immobilization stress and control mice was obtained from GEO (accession: GSE103976). DE analysis was performed by comparing cells from stressed and control mice.

*Ximerakis et al., 2019*^79^. scRNA-seq data from whole brains of young (2-3 mo) and old (21-23 mo) mice from the Broad Institute’s Single Cell Portal (accession: SCP263). DE analysis was performed by comparing cells from young and old mice.

### Visualization

Throughout the manuscript, box plots show the median (horizontal line), interquartile range (hinges) and smallest and largest values no more than 1.5 times the interquartile range (whiskers), and error bars show the standard deviation.

## Supporting information

Supplementary Software 1

## Code availability

We provide an R package, Libra, implementing all methods for DE analysis discussed in this study within a consistent interface. The Libra package is available from GitHub (https://github.com/neurorestore/Libra) and as **Supplementary Software 1**.

## Acknowledgements

We thank L. Adlung, I. Amit, D. Anderson, C. Antelope, D. Arneson, D. Avey, M. Basiri, E. Brenner, G. Chew, M. Co, E. Der, A. Haber, K. Hashimoto, D. Kim, G. Konopka, A. Misharin, R. Mitra, J. Polo, M. Reyes, T. Quertermous, R. Wirka, and O. Yilmaz for providing data and/or cell type annotations. This work was supported by a Consolidator Grant from the European Research Council [ERC-2015-CoG HOW2WALKAGAIN 682999] (to G.C.), the Swiss National Science Foundation (to G.C.; subside 310030_192558), and the Wings for Life Spinal Cord Research Foundation (to M.A.S.). This work was also supported in part by the Intramural Research Program of the NIH, NINDS (to K.J.E.M. and A.L.). Computational resources that supported this work were provided by the Swiss National Supercomputing Center, WestGrid, Compute Canada, and Advanced Research Computing at the University of British Columbia. M.A.S. acknowledges support from the Canadian Institutes of Health Research (CIHR) (Vanier Canada Graduate Scholarship, Michael Smith Foreign Study Supplement), a Vancouver Coastal Health-CIHR-UBC MD/PhD Studentship, and an IUBMB Wood-Whelan Fellowship. J.W.S. is supported by a CIHR Banting Postdoctoral fellowship and a Marie Sklodowska-Curie Individual Fellowship (no. 842578). M.A.A. is supported by a SNF Ambizione fellowship (PZ00P3_185728) and Wings for Life. G.L.M. was supported by the CZI seed network grant HCA3-0000000081 and Swiss National Science Foundation grants CRSK-3_190495 and PZ00P3_193445.

---

**Supplementary Fig. 1.**
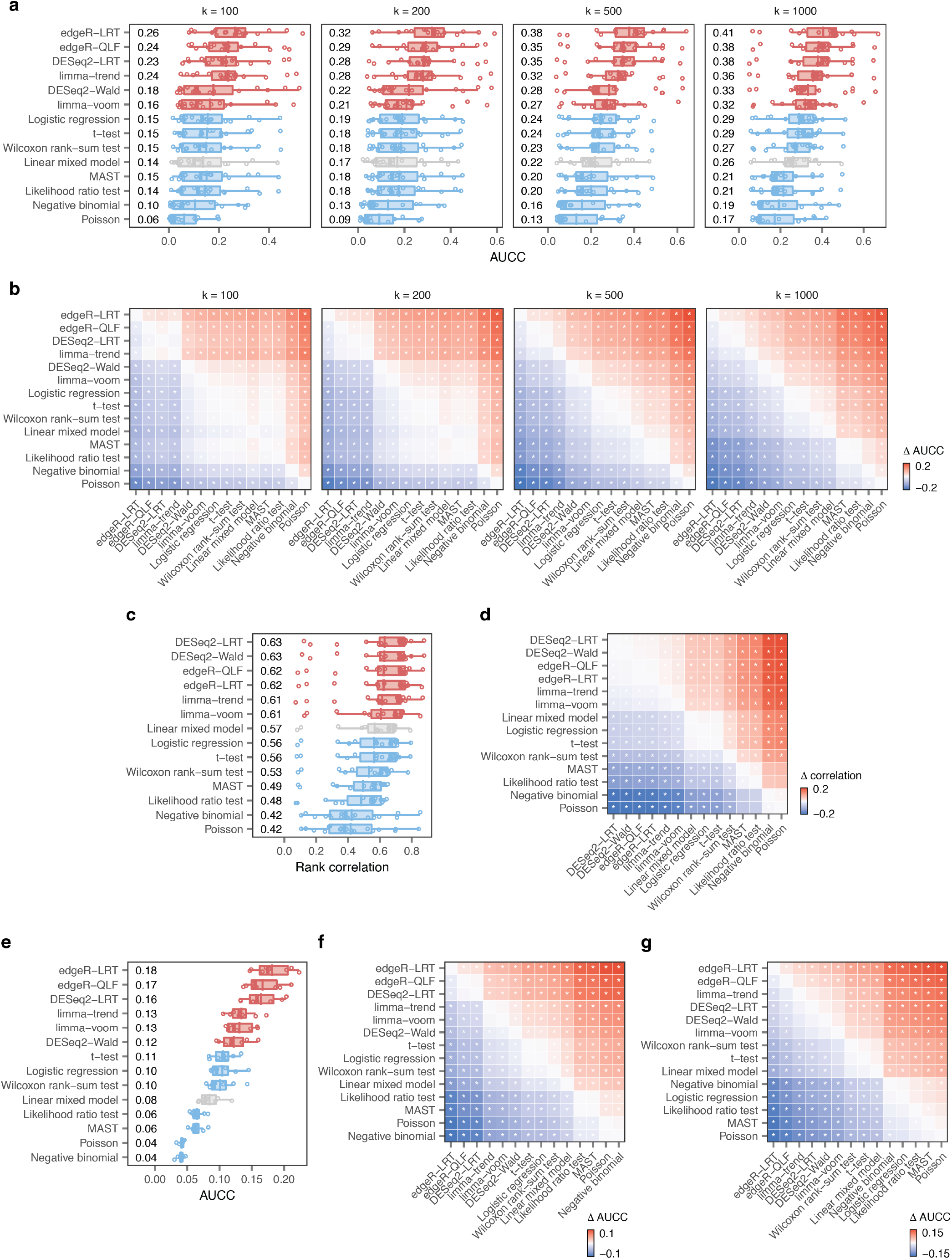
A systematic benchmark of differential expression in single-cell transcriptomics. **a,** Impact of varying the parameter *k* on the AUCC in the eighteen ground-truth datasets, as shown in **Fig. 1c** with *k* = 500. **b,** Impact of varying the parameter k on the ΔAUCC in the eighteen ground-truth datasets, as shown in **Fig. 1d** with k = 500. **c,** Transcriptome-wide rank correlation between single-cell and bulk RNA-seq in the eighteen ground-truth datasets shown in **a**. **d,** Mean difference in the transcriptome-wide rank correlation (Δcorrelation) between the fourteen DE methods shown in **c**. Asterisks indicate comparisons with a two-tailed t-test p-value less than 0.05. **e,** AUCC in eight scRNA-seq datasets with matching bulk proteomics data^14^. **f,** Mean ΔAUCC between the fourteen DE methods shown in **e**. Asterisks indicate comparisons with a two-tailed t-test p-value less than 0.05. **g,** Mean ΔAUCC of GO term enrichment between the fourteen DE methods shown in **Fig. 1e**. Asterisks indicate comparisons with a two-tailed t-test p-value less than 0.05.

**Supplementary Fig. 2.**
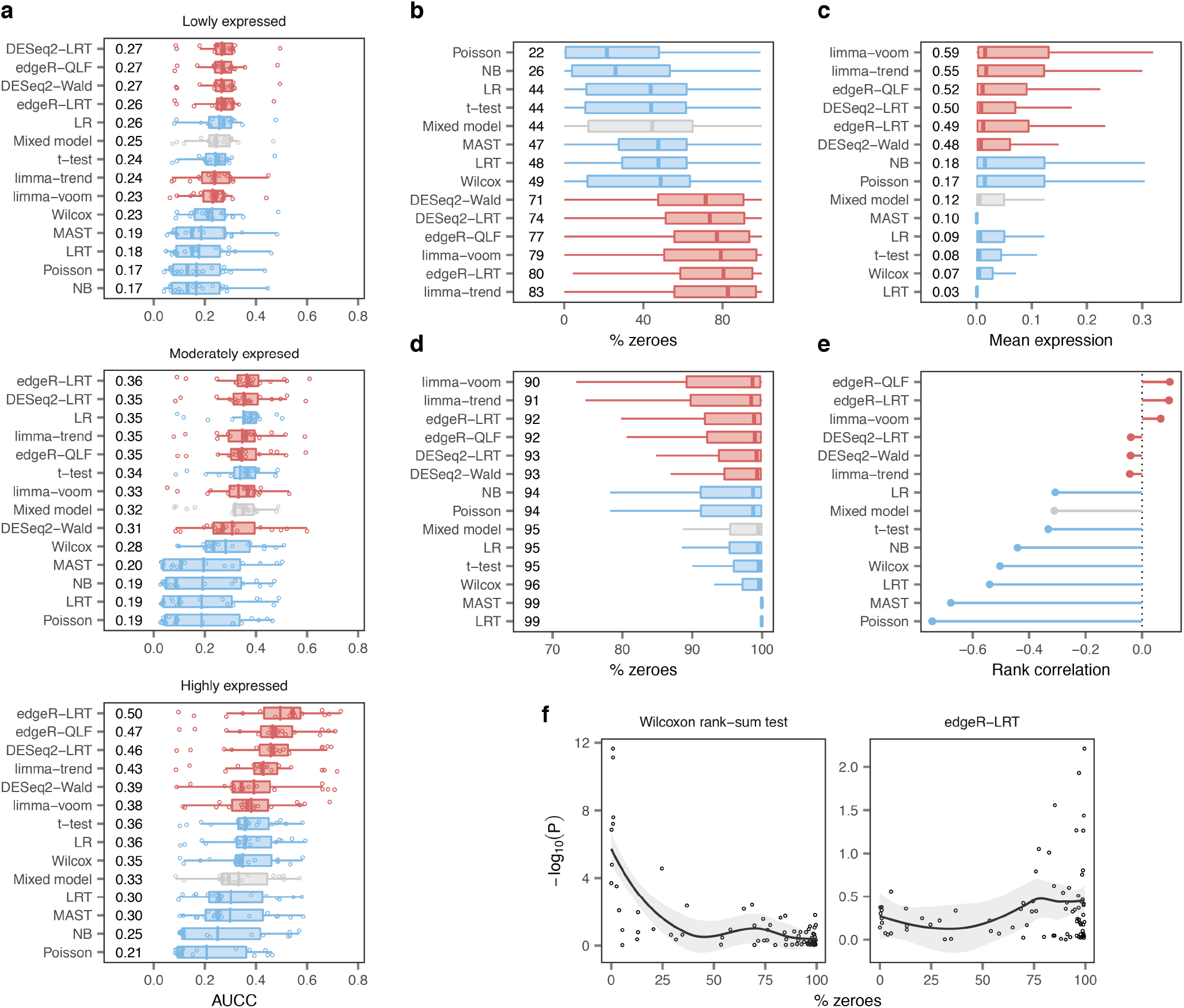
Single-cell DE methods are biased towards highly expressed genes. **a,** AUCCs across eighteen ground-truth datasets after dividing the transcriptome into terciles of lowly (top), moderately (middle), or highly (bottom) expressed genes, as shown in **Fig. 2b**. **b,** Mean proportion of zero gene expression measurements for the 100 top-ranked false-positive genes from each DE method. **c,** Mean expression levels of the 100 top-ranked false-negative genes from each DE method. **d,** Mean proportion of zero gene expression measurements for the 100 top-ranked false-negative genes from each DE method. **e,** Spearman correlation between the mean proportion of zero gene expression measurements for 80 ERCC spike-ins expressed in at least three cells and the −log_10_ p-value of differential expression assigned by each DE method. **f,** Scatterplots of mean proportion of zero gene expression measurements vs. −log_10_ p-value for exemplary single-cell and pseudobulk DE methods.

**Supplementary Fig. 3.**
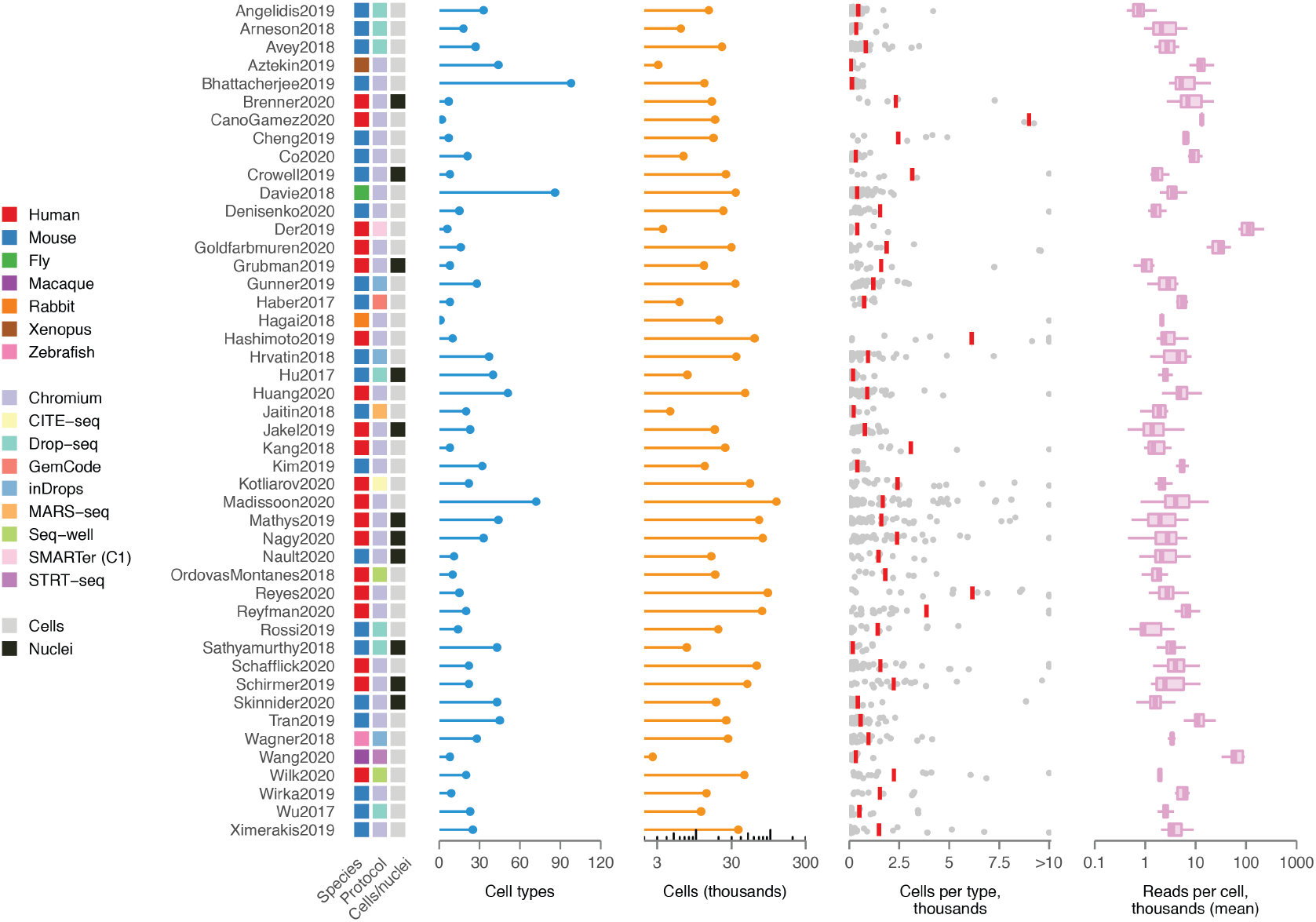
Overview of single-cell transcriptomics datasets. Overview of *n* = 46 published scRNA-seq datasets comparing two or more experimental conditions, used to systematically confirm the universality of the trends observed in analyses of individual datasets. Left, heatmap indicating the species of origin, the sequencing protocol, and whether cells or nuclei were sequenced. Right, properties of each dataset, including the total number of cell types identified in the original studies; the total number of cells sequenced; the number of cells per type (red bars indicate mean); and the mean number of reads for cells of each type.

**Supplementary Fig. 4.**
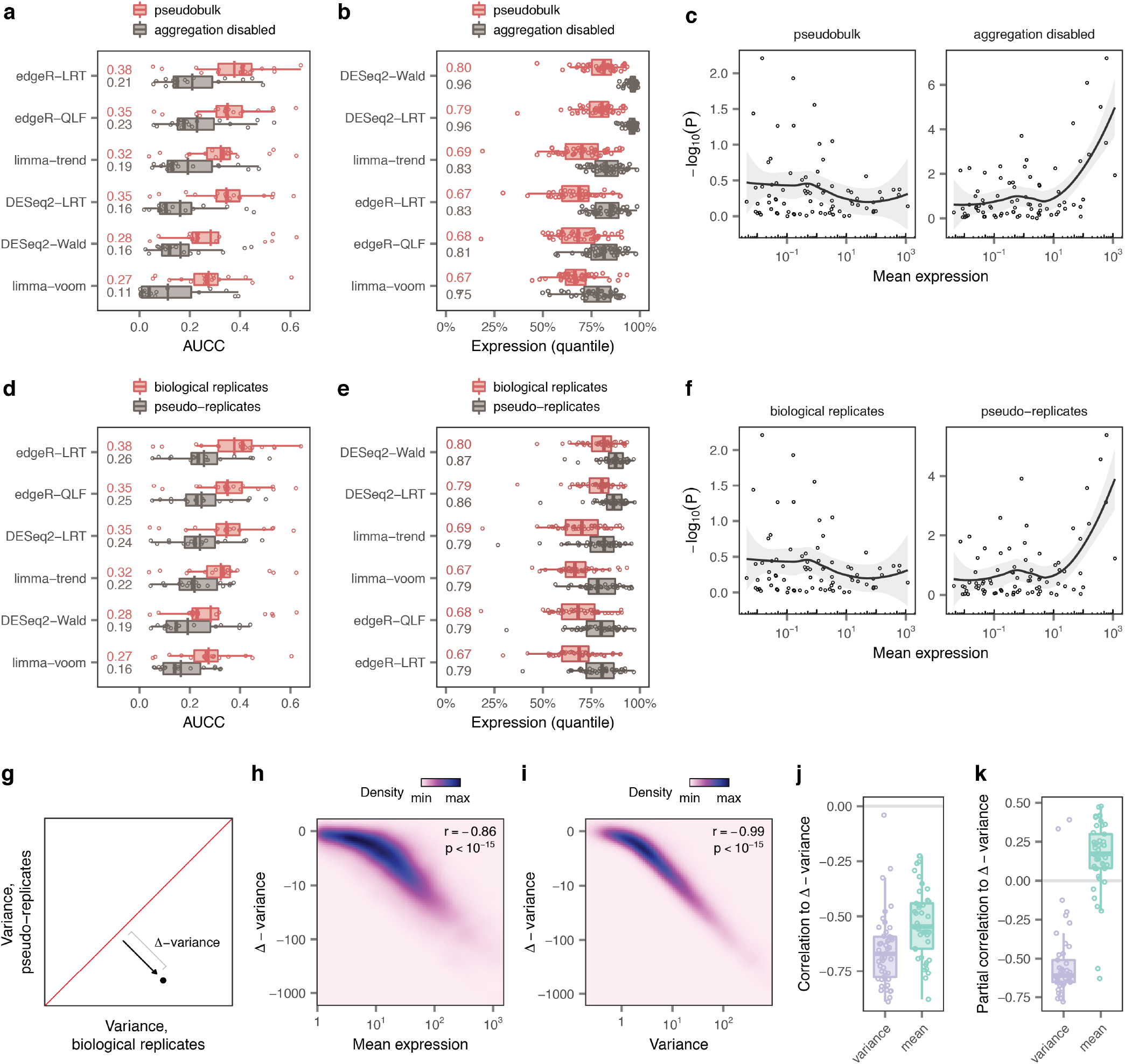
DE analysis in single-cell data must account for biological replicates. **a,** AUCC of the six pseudobulk methods applied to pseudobulks or individual cells in the eighteen ground-truth datasets. **b,** Mean expression levels of the 200 top-ranked genes from six pseudobulk methods applied to pseudobulks or individual cells in a collection of 46 scRNA-seq datasets. **c,** Scatterplots of mean ERCC expression vs. −log10 p-value for an exemplary pseudobulk method, edgeR-LRT, applied to pseudobulks (left) or individual cells (right). **d,** AUCC of the six pseudobulk methods applied to pseudobulks or pseudo-replicates in the eighteen ground-truth datasets. **e,** Mean expression levels of the 200 top-ranked genes from six pseudobulk methods applied to pseudobulks or pseudo-replicates in a collection of 46 scRNA-seq datasets. **f,** Scatterplots of mean ERCC expression vs. −log_10_ p-value for an exemplary pseudobulk method, edgeR-LRT, applied to pseudobulks (left) or pseudoreplicates (right). **g,** Schematic illustrating the calculation of the Δ-variance between biological replicates and pseudo-replicates. **h,** Correlation between mean expression and Δ-variance for 10,448 genes with mean expression ≥ 1 CPM in the dataset of mouse bone marrow mononuclear cells stimulated with poly-I:C. Mean expression is strongly correlated with Δ-variance, such that the variance of highly expressed genes is disproportionately underestimated when ignoring information about biological replicates. **i,** Correlation between expression variance and Δ-variance for 10,448 genes with mean expression ≥ 1 CPM in the dataset of mouse bone marrow mononuclear cells stimulated with poly-I:C. Variance is even more strongly correlated with Δ-variance than mean expression, such that the most variable genes are disproportionately underestimated when ignoring information about biological replicate. **j,** Correlation between mean expression levels or expression variance and Δ-variance in 46 scRNA-seq datasets. Variance is even more strongly correlated with Δ-variance than mean expression across a large compendium of datasets, corroborating the trends shown in **h-i**. **k,** Partial correlation between mean expression and Δ-variance, controlling for variance, or between variance and Δ-variance, controlling for mean expression. The variance of gene expression is the primary determinant of Δ-variance, implying that failing to account for biological replicates introduces a bias towards highly expressed genes because these genes are also more variable.

**Supplementary Fig. 5.**
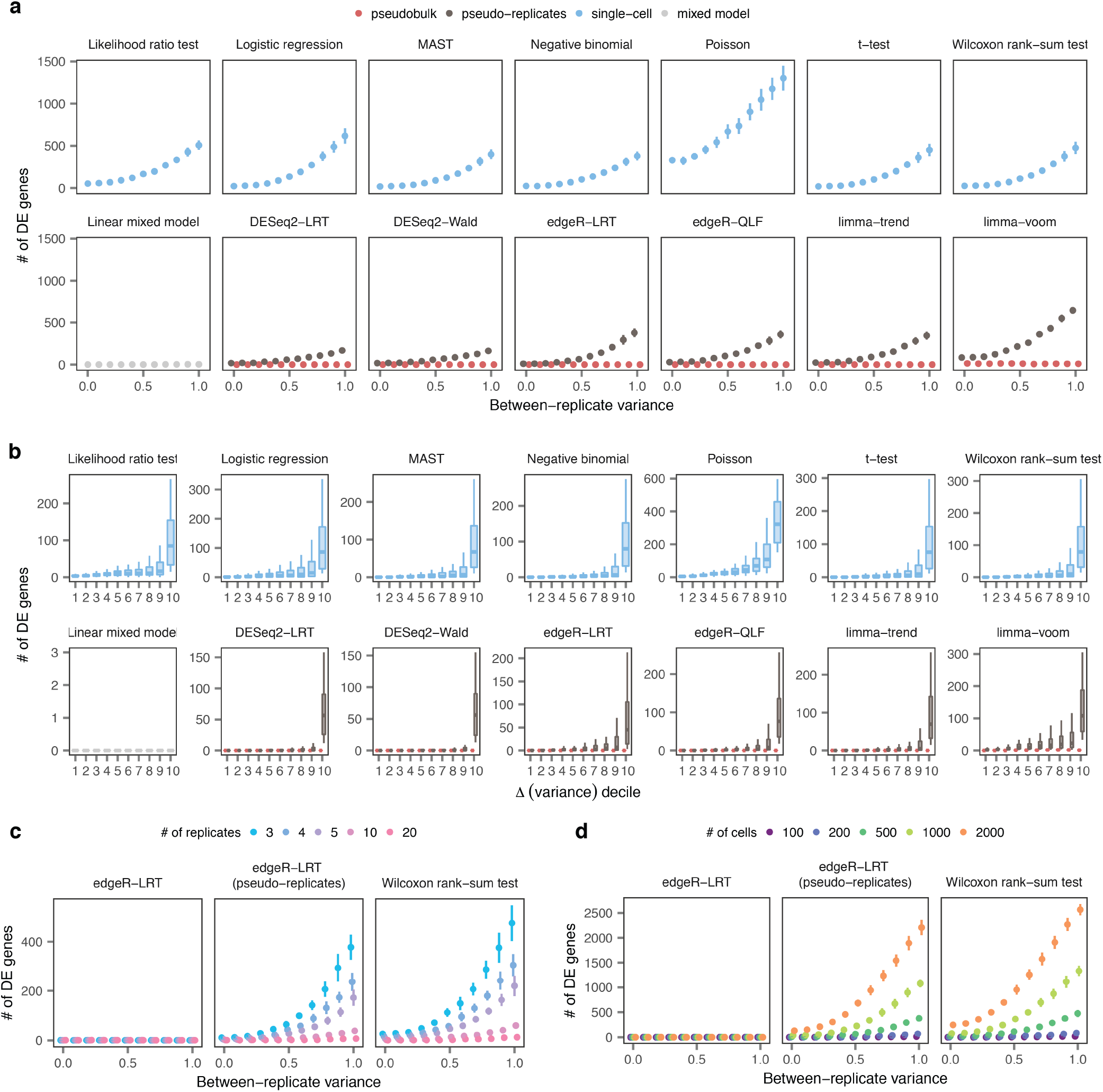
Simulation studies expose false discoveries in single-cell DE. **a,** Number of DE genes detected in stimulation experiments with varying degrees of heterogeneity between replicates by all DE methods. **b,** Number of DE genes detected by the tests shown in a for genes divided into deciles by the magnitude of the change in variance between biological replicates and pseudo-replicates (Δ-variance). **c,** Number of DE genes detected by a representative single-cell DE method, a representative pseudobulk method, and the same pseudobulk method applied to pseudo-replicates, when varying the total number of replicates in the simulated dataset. **d,** Number of DE genes detected by a representative single-cell DE method, a representative pseudobulk method, and the same pseudobulk method applied to pseudo-replicates, when varying the total number of cells in the simulated dataset.

**Supplementary Fig. 6.**
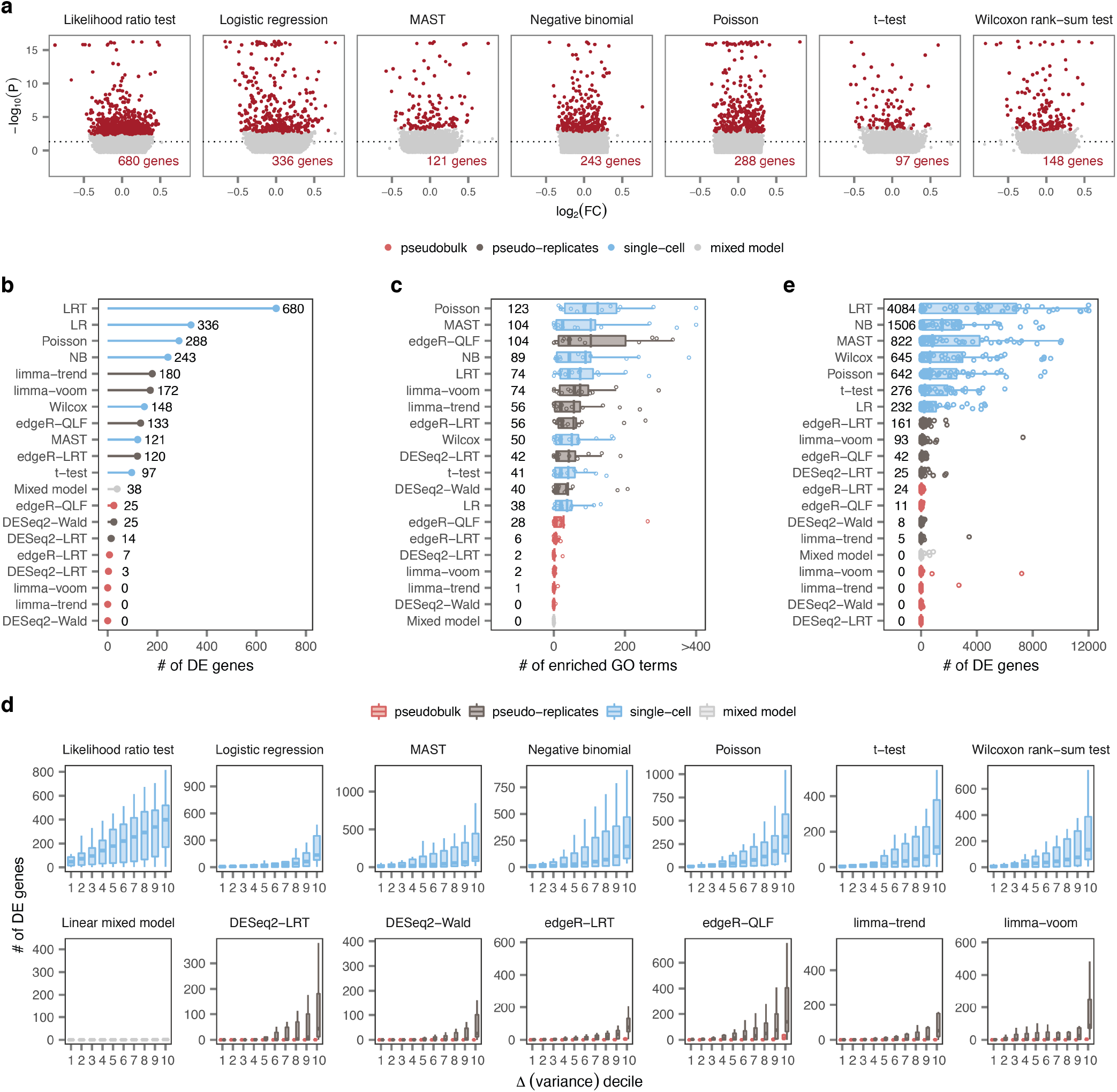
False discoveries in single-cell and spatial transcriptomics data. **a,** Volcano plots showing DE between T cells from random groups of unstimulated controls drawn from Kang et al.^5^ using seven single-cell DE methods. **b,** Number of DE genes detected by all DE methods in unstimulated T cells. **c,** Number of GO terms enriched at 5% FDR among DE genes identified in comparisons of random groups of unstimulated controls from fourteen scRNA-seq studies with at least six control samples. **d,** Number of DE genes in comparisons of random groups of unstimulated controls from fourteen scRNA-seq studies with at least six control samples, as shown in **Fig. 4e**, for genes divided into deciles by the magnitude of the change in variance between biological replicates and pseudo-replicates (Δ-variance). **e,** Number of DE genes detected by all DE methods within spinal cord regions from control mice profiled by spatial transcriptomics^24^.

**Supplementary Fig. 7.**
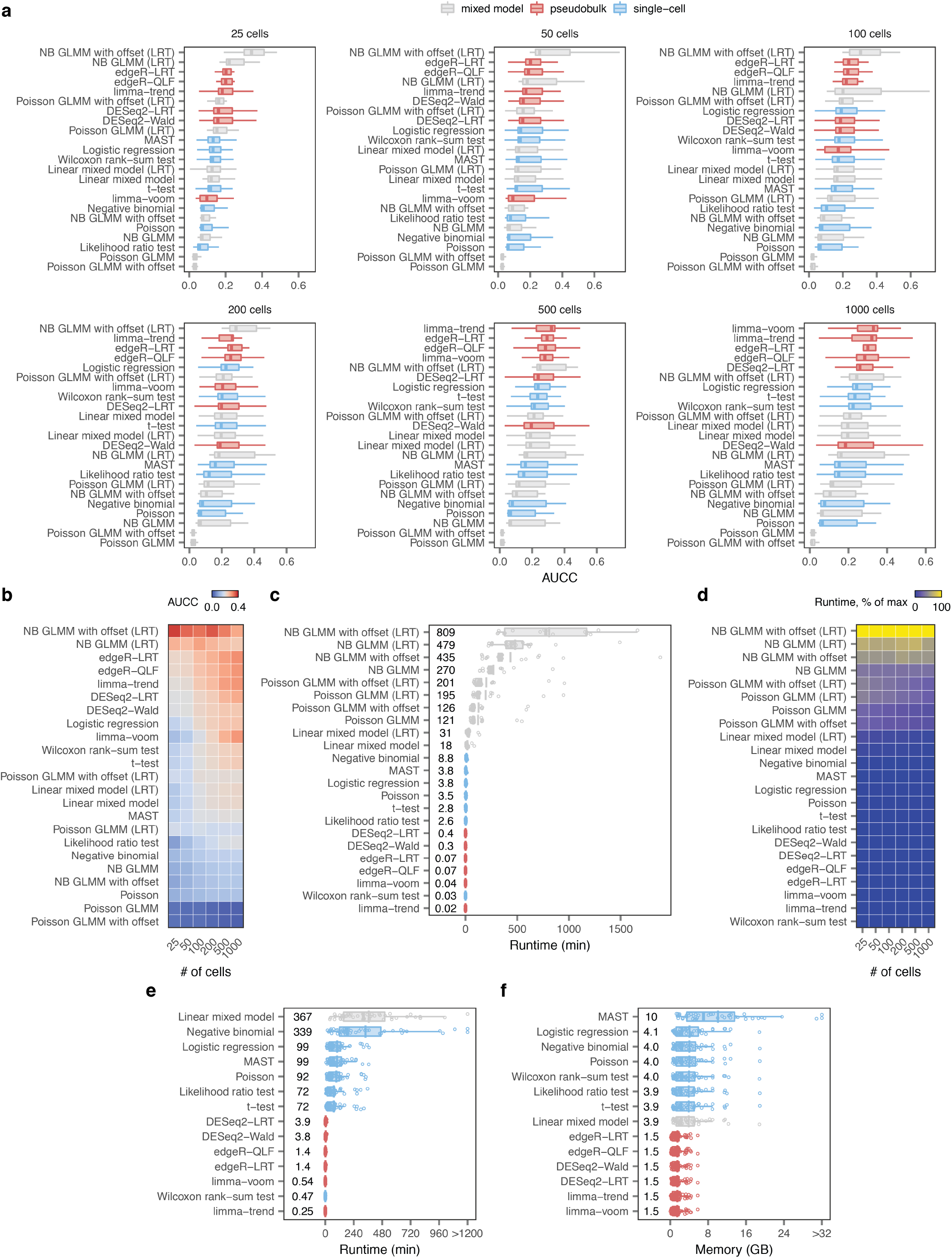
Single-cell DE analysis with generalized linear mixed models. **a,** AUCC for ten different generalized linear mixed models (GLMMs), varying in the choice of link function (identity, Poisson, or negative binomial, NB); method used to evaluate statistical significance (Wald test or likelihood ratio test, LRT), and presence of an offset term, in samples of between 25 and 1,000 cells from the eighteen ground-truth datasets shown in **Fig. 1c**, and compared to the fourteen DE methods shown in the same panel. **b,** As in **a,** but showing the mean AUCC as a function of the number of cells sampled for each DE method. **c,** Runtime in minutes for the ten GLMMs shown in a in samples of 1,000 cells. The top-performing GLMM required a mean of 13.5 h per cell type to perform DE analysis. **d,** Runtime of the ten GLMMs and the fourteen DE methods shown in **Fig. 1c**, shown as a percentage of the maximum runtime, as a function of the number of cells sampled. **e,** Runtime in minutes of the fourteen DE methods shown in **Fig. 1c** across 46 scRNA-seq datasets. **f,** Maximum memory required in gigabytes by the fourteen DE methods shown in **Fig. 1c** across 46 scRNA-seq datasets.

