## Supplementary Software 1 for "Confronting false discoveries in single-cell differential expression": check_inputs.R

```
#' Check inputs
#' 
#' Check inputs prior to running Libra functions
#'
#' 
#' @param input a single-cell matrix to be converted, with features (genes) in rows
#'   and cells in columns. Alternatively, a \code{Seurat}, \code{monocole3}, or 
#'   or \code{SingleCellExperiment} object can be directly input.
#' @param meta the accompanying meta data whereby the rownames match the column
#'   names of \code{input}. 
#' @param replicate_col the vector in \code{meta} containing the replicate 
#'   information. Defaults to \code{replicate}.
#' @param cell_type_col the vector in \code{meta} containing the cell type 
#'   information. Defaults to \code{cell_type}.
#' @param label_col the vector in \code{meta} containing the experimental
#'   label. Defaults to \code{label}. 
#' @param min_cells the minimum number of cells in a cell type to retain it.
#'   Defaults to \code{3}.
#' @param min_reps the minimum number of replicates in a cell type to retain it.
#'   Defaults to \code{2}.
#' @param min_features the minimum number of counts for a gene to retain it.
#'   Defaults to \code{0}   
#' @return a cleaned up expression matrix and meta data object
#'  
#' @importFrom dplyr %>% rename_ n_distinct mutate_at vars
#' @importFrom tester is_numeric_matrix is_numeric_dataframe
#' @importFrom methods is
#' 
check_inputs = function(input, 
                        meta = NULL, 
                        replicate_col = 'replicate',
                        cell_type_col = 'cell_type',
                        label_col = 'label') {
  # extract cell types and label from metadata
  if ("Seurat" %in% class(input)) {
    # confirm Seurat is installed
    if (!requireNamespace("Seurat", quietly = TRUE)) {
      stop("install \"Seurat\" R package for Augur compatibility with ",
           "input Seurat object", call. = FALSE)
    }
    meta = input@meta.data %>%
      droplevels()
    if (!is.null(replicate_col))
      replicates = as.character(meta[[replicate_col]])
    labels = as.character(meta[[label_col]])
    cell_types = as.character(meta[[cell_type_col]])
    expr = Seurat::GetAssayData(input, slot = 'counts')
  } else if ("cell_data_set" %in% class(input)) {
    # confirm monocle3 is installed
    if (!requireNamespace("monocle3", quietly = TRUE)) {
      stop("install \"monocle3\" R package for Augur compatibility with ",
           "input monocle3 object", call. = FALSE)
    }
    meta = monocle3::pData(input) %>%
      droplevels() %>%
      as.data.frame()
    if (!is.null(replicate_col))
      replicates = as.character(meta[[replicate_col]])
    labels = as.character(meta[[label_col]])
    cell_types = as.character(meta[[cell_type_col]])
    expr = monocle3::exprs(input)
  } else if ("SingleCellExperiment" %in% class(input)){
    # confirm SingleCellExperiment is installed
    if (!requireNamespace("SingleCellExperiment", quietly = TRUE)) {
      stop("install \"SingleCellExperiment\" R package for Augur ",
           "compatibility with input SingleCellExperiment object",
           call. = FALSE)
    }
    meta = SummarizedExperiment::colData(input) %>%
      droplevels() %>%
      as.data.frame()
    if (!is.null(replicate_col))
      replicates = as.character(meta[[replicate_col]])
    labels = as.character(meta[[label_col]])
    cell_types = as.character(meta[[cell_type_col]])
    expr = SummarizedExperiment::assay(input)
  } else {
    # check if input is sparse matrix or numberic matrix/df
    valid_input = is(input, 'sparseMatrix') ||
      is_numeric_matrix(input) ||
      is_numeric_dataframe(input)
    if (!valid_input)
      stop("input must be Seurat, monocle, sparse matrix, numeric matrix, or ",
           "numeric data frame")
    if (is.null(meta)) 
      stop("input matrix must be accompanied by a metadata table")
    expr = input
    if (!is.null(replicate_col))
      replicates = as.character(meta[[replicate_col]])
    labels = as.character(meta[[label_col]])
    cell_types = as.character(meta[[cell_type_col]])
  }
  
  # check dimensions are non-zero
  if (length(dim(expr)) != 2 || !all(dim(expr) > 0)) {
    stop("expression matrix has at least one dimension of size zero")
  }
  
  # check dimensions match
  n_cells1 = nrow(meta)
  n_cells2 = ncol(expr)
  if (n_cells1 != n_cells2) {
    stop("number of cells in metadata (", n_cells1, ") does not match number ",
         "of cells in expression (", n_cells2, ")")
  }
  
  # check at least two labels
  if (n_distinct(labels) == 1) {
    stop("only one label provided: ", unique(labels))
  }
  
  # check for missing labels or cell types
  if (any(is.na(labels))) {
    stop("labels contain ", sum(is.na(labels)), "missing values")
  }
  if (any(is.na(cell_types))) {
    stop("cell types contain ", sum(is.na(cell_types)), "missing values")
  }
  if (!is.null(replicate_col) && any(is.na(replicates))) {
    stop("replicates contain ", sum(is.na(replicates)), "missing values")
  }
  
  # check for missing replicates
  if (!is.null(replicate_col) && is.null(replicates)) {
    stop("metadata does not contain replicate information")
  }
  
  # remove missing values
  missing = is.na(expr)
  if (any(missing)) {
    stop("matrix contains ", sum(missing), "missing values")
  }
  
  # clean up the meta data
  if (!is.null(replicate_col)) {
    meta %<>%
      mutate(cell_barcode = rownames(meta),
             replicate = meta[[replicate_col]],
             cell_type = meta[[cell_type_col]],
             label = meta[[label_col]]) %>%
      mutate_at(vars(replicate, cell_type, label), as.factor)
  } else {
    meta %<>%
      mutate(cell_barcode = rownames(meta),
             cell_type = meta[[cell_type_col]],
             label = meta[[label_col]]) %>%
      mutate_at(vars(cell_type, label), as.factor)
  }
  
  # make sure we only have 2 labels
  if (n_distinct(meta$label) != 2) {
    stop("Please only include 2 distinct experimental labels")
  }
  # make sure meta contains row names and is a data frame
  rownames(meta) = colnames(expr)
  meta = as.data.frame(meta)
  to_return = list(
    expr = expr,
    meta = meta
  )
  return(to_return)
}
```
