## Supplementary Software 1 for "Confronting false discoveries in single-cell differential expression": hagai_toy.R

```
#' A toy single-cell RNA-seq dataset derived from Hagai et al., 2018 Nature
#' 
#' A toy simulated scRNA-seq dataset, used to demonstrate Libra 
#' The dataset contains one population of cells:
#' bone marrow derived mononuclear phagocytes.
#' Each label (unst and lps4) contain three replicates, each with 100 cells
#' For speed we have included only 100 genes.
#' The data is provided as a Seurat object. 
#' 
#' @docType data
#' @usage data(hagai_toy)
#' @format a Seurat object with dimensions 100 genes x 600 cells
"hagai_toy"
```
