## Supplementary Software 1 for "Confronting false discoveries in single-cell differential expression": mixedmodel_de.R

```
#' Run mixed model differential expression
#' 
#' Run differential expression, accounting for biological replicates in
#' single cell data. Options include pseudobulk or mixed model approaches
#' 
#' @param input a single-cell matrix to be converted, with features (genes) in rows
#'   and cells in columns. Alternatively, a \code{Seurat}, \code{monocle3}, or 
#'   or \code{SingleCellExperiment} object can be directly input.
#' @param meta the accompanying meta data whereby the rownames match the column
#'   names of \code{input}. 
#' @param replicate_col the vector in \code{meta} containing the replicate 
#'   information. Defaults to \code{replicate}.
#' @param cell_type_col the vector in \code{meta} containing the cell type 
#'   information. Defaults to \code{cell_type}.
#' @param label_col the vector in \code{meta} containing the experimental
#'   label. Defaults to \code{label}. 
#' @param min_cells the minimum number of cells in a cell type to retain it.
#'   Defaults to \code{3}.
#' @param min_reps the minimum number of replicates in a cell type to retain it.
#'   Defaults to \code{2}.
#' @param min_features the minimum number of counts for a gene to retain it.
#'   Defaults to \code{0}   
#' @param de_method the mixed model type to use. Defaults to negbinom.
#' @param de_type the specific mixed model test to use. Defaults to LRT.
#' @param n_threads number of threads to use for parallelization.
#' @param show_progress show a progress bar. Defaults to T
#' @return a data frame containing differential expression results.
#'  
#' @importFrom pbmcapply pbmclapply
#' @importFrom parallel mclapply
#' @importFrom dplyr count group_by filter pull mutate
#' @importFrom magrittr extract set_rownames
#' @importFrom Matrix colSums rowMeans
#' @importFrom lmerTest lmer
#' @importFrom lme4 lmerControl .makeCC
#' @importFrom glmmTMB glmmTMB nbinom1
#' @importFrom blme bglmer
#' @importFrom stats coef anova as.formula p.adjust model.matrix
#' @importFrom Seurat CreateSeuratObject NormalizeData Idents GetAssayData
#'   WhichCells
#' @importFrom methods new
#' 

mixedmodel_de = function(
  input,
  meta = NULL,
  replicate_col = 'replicate',
  cell_type_col = 'cell_type',
  label_col = 'label',
  latent_vars = NULL,
  min_cells = 3,
  min_features = 0,
  de_family = 'mixedmodel',
  de_method = 'negbinom',
  de_type = 'LRT',
  n_threads = 2,
  show_progress = T
) {
  # check the arguments
  if (!de_method %in% c("negbinom", "linear", "poisson",
                        "negbinom_offset", "poisson_offset"))
    stop("Please select one of: negbinom, linear, poisson,
         negbinom_offset or poisson_offset as the model")
  if(!de_type %in% c("LRT", "Wald"))
    stop("Please select one of: LRT, Wald as the statistical test")
  
  # first, make sure inputs are correct
  inputs = check_inputs(
    input, 
    meta,
    replicate_col = replicate_col,
    cell_type_col = cell_type_col,
    label_col = label_col
    )
  expr = inputs$expr
  meta = inputs$meta
  
  # define the formula to use
  fmla <- paste(
    "GENE ~",
    paste(label_col, latent_vars, collapse = "+")
  )
  
  if (grepl("offset", de_method)) {
    use_offset = T
    # tag the de method
    de_tag = de_method
    de_method = gsub("_offset", "", de_method)
  } else {
    use_offset = F
    # tag the de method
    de_tag = de_method
  }
  
  # add in the offset if required
  if (use_offset == F) {
    fmla = as.formula(paste(fmla, "+", "(1|replicate)"))
    message("..using formula: ",
            paste0("GENE ~ ", paste(label_col, latent_vars, collapse = "+")),
            " with random term (1|replicate)"
    )
  } else if (use_offset == T) {
    fmla = as.formula(paste(fmla, "+ offset(log(total_counts)) +", "(1|replicate)"))
    message("..using formula: ",
            paste0("GENE ~ ", paste(label_col, latent_vars, collapse = "+")),
            "with a total counts correction term offset(log(total_counts)) and",
            " with random term (1|replicate)"
    )
  }
  
  # check if showing progress or not
  if (show_progress == T) {
    apply_fun = pbmclapply
  } else {
    apply_fun = mclapply
  }
  
  # identify cell types
  cell_types = unique(meta$cell_type)
  
  # keep only cell types with enough cells
  keep = meta %>%
    dplyr::count(cell_type, label) %>%
    group_by(cell_type) %>%
    filter(all(n >= min_cells)) %>%
    pull(cell_type) %>%
    unique()
  cell_types = cell_types[cell_types %in% keep]
  
  # check minimum features
  keep = rowSums(expr) > min_features
  expr = expr[keep,]
  
  for (cell_type in cell_types) {
    # get the subset matrix
    meta0 = meta %>% filter(cell_type == !!cell_type)
    expr0 = expr %>% extract(, meta0$cell_barcode)
    # define genes to use
    genes = rownames(expr0)
    result = apply_fun(genes, mc.cores = n_threads, function(x) {
      test_dat = data.frame(
        GENE = expr[x,],
        label = meta0$label,
        replicate = meta0$replicate
      )
      if (use_offset == T) {
        test_dat %<>% mutate(total_counts = as.numeric(colSums(expr0)+1))
      }
      # define coefficient
      coef = paste0("label", levels(test_dat$label)[2])
      DE = tryCatch({
        switch(de_method,
               linear = {
                 if (de_type == 'LRT') {
                   null_fmla = as.formula(gsub("label \\+ ", " ", deparse(fmla)))
                   mod1 = lmer(fmla, test_dat, REML = TRUE, control = lmerControl(
                     check.conv.singular = .makeCC(action = "ignore", tol = 1e-4)))
                   mod2 = lmer(null_fmla, test_dat, REML = TRUE, control = lmerControl(
                     check.conv.singular = .makeCC(action = "ignore", tol = 1e-4)))
                   lrt = anova(mod1, mod2, refit = F)
                   p_val = lrt$`Pr(>Chisq)`[2]
                   test_statistic = lrt$Chisq[2]
                 } else if (de_type == 'Wald') {
                   tab <- coef(summary(
                     lmer(fmla, test_dat, REML = TRUE, control = lmerControl(
                       check.conv.singular = .makeCC(action = "ignore", tol = 1e-4)))
                   ))
                   p_val <- tab[coef,5]
                   test_statistic <- tab[coef,4]
                 }
                 c(p_val, test_statistic)
               },
               negbinom = {
                 if (de_type == 'LRT') {
                   null_fmla = as.formula(gsub("label \\+ ", " ", deparse(fmla)))
                   mod1 = glmmTMB(fmla, test_dat, family = nbinom1, REML = FALSE)
                   mod2 = glmmTMB(null_fmla, test_dat, family = nbinom1, REML = FALSE)
                   lrt = anova(mod1, mod2, refit = F)
                   p_val = lrt$`Pr(>Chisq)`[2]
                   test_statistic = lrt$Chisq[2]
                 } else if (de_type == "Wald") {
                   tab <- coef(summary(
                     glmmTMB(fmla, test_dat, family = nbinom1, REML = FALSE)
                   ))[[1]]
                   p_val <- tab[coef,4]
                   test_statistic <- tab[coef,3]
                 }
                 c(p_val, test_statistic)
               },
               poisson = {
                 if (de_type == 'LRT') {
                   null_fmla = as.formula(gsub("label \\+ ", " ", deparse(fmla)))
                   mod1 = bglmer(fmla, test_dat, family = 'poisson')
                   mod2 = bglmer(null_fmla, test_dat, family = 'poisson')
                   lrt = anova(mod1, mod2, refit = F)
                   p_val = lrt$`Pr(>Chisq)`[2]
                   test_statistic = lrt$Chisq[2]
                 } else if (de_type == 'Wald') {
                   tab <- coef(summary(
                     bglmer(fmla, test_dat, family = 'poisson')
                   ))
                   p_val <- tab[coef, 4]
                   test_statistic <- tab[coef, 3]
                 }
                 c(p_val, test_statistic)
               }
        )
      }, error = function(e) c(NA_real_, NA_real_)
      )
    })
    # grab the logFC for this cell type, using tp10K
    logFC = tryCatch({
      sc0 = CreateSeuratObject(
        expr0, 
        meta = meta0 %>% set_rownames(.$cell_barcode)) %>%
        NormalizeData()
      Idents(sc0) = sc0$label
      mat = GetAssayData(sc0, slot = 'data')
      levels = levels(meta0$label)
      if (is.null(levels)) {
        levels = unique(meta0$label)
      }
      cells1 = WhichCells(sc0, idents = levels[1])
      cells2 = WhichCells(sc0, idents = levels[2])
      data1 = log(rowMeans(mat[, cells1, drop = F] + 1))
      data2 = log(rowMeans(mat[, cells2, drop = F] + 1))
      out = as.numeric(data2 - data1)
    }, error = function(e) { return(NA_real_) })
    # format the results properly
    vec = unlist(result)
    p_val = vec[seq(1,length(vec),2)]
    test_statistic = vec[seq(2,length(vec),2)]
    return(data.frame(p_val = p_val, 
                      test_statistic = test_statistic,
                      gene = rownames(expr)) %>%
             set_rownames(rownames(expr)) %>%
             mutate(p_val_adj = p.adjust(p_val, method = 'BH'),
                    cell_type = !!cell_type,
                    logFC = logFC,
                    de_family = 'pseudobulk',
                    de_method = de_tag,
                    de_type = de_type))
  }
}
  
  
```
