## Supplementary Software 1 for "Confronting false discoveries in single-cell differential expression": pseudobulk_de.R

```
#' Run pseudobulk differential expression methods
#' 
#' Run pseudobulk differential expression methods on single-cell data
#' 
#' @param input a single-cell matrix to be converted, with features (genes) in rows
#'   and cells in columns. Alternatively, a \code{Seurat}, \code{monocle3}, or 
#'   or \code{SingleCellExperiment} object can be directly input.
#' @param meta the accompanying meta data whereby the rownames match the column
#'   names of \code{input}.
#' @param replicate_col the vector in \code{meta} containing the replicate 
#'   information. Defaults to \code{replicate}.
#' @param cell_type_col the vector in \code{meta} containing the cell type 
#'   information. Defaults to \code{cell_type}.
#' @param label_col the vector in \code{meta} containing the experimental
#'   label. Defaults to \code{label}. 
#' @param min_cells the minimum number of cells in a cell type to retain it.
#'   Defaults to \code{3}.
#' @param min_reps the minimum number of replicates in a cell type to retain it.
#'   Defaults to \code{2}.
#' @param min_features the minimum number of counts for a gene to retain it.
#'   Defaults to \code{0}   
#' @param de_method the specific differential expression testing method to use.
#'   Defaults to edgeR.
#' @param de_type the specific parameter of the differential expression testing
#'   method. Defaults to LRT for edgeR, LRT for DESeq2, and trend for limma.
#' @return a data frame containing differential expression results.
#'  
#' @importFrom magrittr %<>%
#' @importFrom tibble rownames_to_column
#' @importFrom dplyr %>% mutate n_distinct
#' @importFrom edgeR DGEList calcNormFactors estimateDisp glmQLFit glmQLFTest
#'   glmFit glmLRT topTags cpm
#' @importFrom DESeq2 DESeqDataSetFromMatrix DESeq results
#' @importFrom limma voom lmFit eBayes topTable
#' @importFrom purrr map 
#' @importFrom stats model.matrix
#' @importFrom methods new
#' 
#' 
pseudobulk_de = function(input, 
                         meta = NULL, 
                         replicate_col = 'replicate',
                         cell_type_col = 'cell_type',
                         label_col = 'label',
                         min_cells = 3,
                         min_reps = 2,
                         min_features = 0,
                         de_family = 'pseudobulk',
                         de_method = 'edgeR',
                         de_type = 'LRT') {
  # check args
  if (de_method == 'limma') {
    # change default type to use
    de_type = 'trend'
  }

  # first, make sure inputs are correct
  inputs = check_inputs(
    input = input,
    meta = meta,
    replicate_col = replicate_col,
    cell_type_col = cell_type_col,
    label_col = label_col)
  expr = inputs$expr
  meta = inputs$meta
  
  # get the pseudobulks list
  pseudobulks = to_pseudobulk(
    input = expr,
    meta = meta,
    replicate_col = replicate_col,
    cell_type_col = cell_type_col,
    label_col = label_col,
    min_cells = min_cells,
    min_reps = min_reps,
    min_features = min_features
  )
  
  #TODO: wrap this in map
  results = map(pseudobulks, function(x) {
    # create targets matrix
    targets = data.frame(group_sample = colnames(x)) %>%
      mutate(group = gsub(".*\\:", "", group_sample))
    ## optionally, carry over factor levels from entire dataset
    if (is.factor(meta$label)) {
      targets$group %<>% factor(levels = levels(meta$label))
    }
    if (n_distinct(targets$group) > 2)
      return(NULL)
    # create design
    design = model.matrix(~ group, data = targets)
    
    DE = switch(de_method,
                edgeR = {
                  tryCatch({
                    y = DGEList(counts = x, group = targets$group) %>%
                      calcNormFactors(method = 'TMM') %>%
                      estimateDisp(design)
                    test = switch(de_type,
                                  QLF = {
                                    fit = glmQLFit(y, design)
                                    test = glmQLFTest(fit, coef = -1)
                                  },
                                  LRT = {
                                    fit = glmFit(y, design = design)
                                    test = glmLRT(fit)
                                  })
                    res = topTags(test, n = Inf) %>%
                      as.data.frame() %>%
                      rownames_to_column('gene') %>%
                      # flag metrics in results
                      mutate(de_family = 'pseudobulk',
                             de_method = de_method,
                             de_type = de_type)
                  }, error = function(e) {
                    message(e)
                    data.frame()
                  })
                },
                DESeq2 = {
                  tryCatch({
                    dds = DESeqDataSetFromMatrix(countData = x,
                                                 colData = targets,
                                                 design = ~ group)
                    dds = switch(de_type,
                                 Wald = {
                                   dds = try(DESeq(dds,
                                                   test = 'Wald',
                                                   fitType = 'parametric',
                                                   sfType = 'poscounts',
                                                   betaPrior = F))
                                 },
                                 LRT = {
                                   dds = try(DESeq(dds,
                                                   test = 'LRT',
                                                   reduced = ~ 1,
                                                   fitType = 'parametric',
                                                   sfType = 'poscounts',
                                                   betaPrior = F))
                                 }
                    )
                    res = results(dds)
                    # write
                    res = as.data.frame(res) %>%
                      mutate(gene = rownames(x)) %>%
                      # flag metrics in results
                      mutate(de_family = 'pseudobulk',
                             de_method = de_method,
                             de_type = de_type)
                  }, error = function(e) {
                    message(e)
                    data.frame()
                  })
                },
                limma = {
                  tryCatch({
                    x = switch(de_type,
                               trend = {
                                 trend_bool = T
                                 dge = DGEList(as.matrix(x), group = targets$group)
                                 dge = calcNormFactors(dge)
                                 x = new("EList")
                                 x$E = cpm(dge, log = TRUE, prior.count = 3)
                                 x
                               },
                               voom = {
                                 counts = all(as.matrix(x) %% 1 == 0)
                                 if (counts) {
                                   trend_bool = F
                                   x = voom(as.matrix(x), design)
                                   x
                                 }
                               })
                    # get fit
                    fit = lmFit(x, design) %>%
                      eBayes(trend = trend_bool, robust = trend_bool)
                    # format the results
                    res = fit %>%
                      # extract all coefs except intercept
                      topTable(number = Inf, coef = -1) %>%
                      rownames_to_column('gene') %>%
                      # flag metrics in results
                      mutate(
                        de_family = 'pseudobulk',
                        de_method = de_method,
                        de_type = de_type)
                  }, error = function(e) {
                    message(e)
                    data.frame()
                  })
                }
    )
  })
  results %<>% bind_rows(.id = 'cell_type')
}
```
