## Supplementary Software 1 for "Confronting false discoveries in single-cell differential expression": run_de.R

```
#' Run differential expression
#' 
#' Perform differential expression on single-cell data. Libra implements a total 
#' of 22 unique differential expression methods that can all be accessed from 
#' one function. These methods encompass traditional single-cell methods as well 
#' as methods accounting for biological replicate including pseudobulk and 
#' mixed model methods. The code for this package has been largely inspired 
#' by the Seurat and Muscat packages. Please see the documentation of these 
#' packages for further information.
#' 
#' @param input a single-cell matrix to be converted, with features (genes) in rows
#'   and cells in columns. Alternatively, a \code{Seurat}, \code{monocle3}, or 
#'   or \code{SingleCellExperiment} object can be directly input.
#' @param meta the accompanying meta data whereby the rownames match the column
#'   names of \code{input}. 
#' @param replicate_col the vector in \code{meta} containing the replicate 
#'   information. Defaults to \code{replicate}.
#' @param cell_type_col the vector in \code{meta} containing the cell type 
#'   information. Defaults to \code{cell_type}.
#' @param label_col the vector in \code{meta} containing the experimental
#'   label. Defaults to \code{label}. 
#' @param min_cells the minimum number of cells in a cell type to retain it.
#'   Defaults to \code{3}.
#' @param min_reps the minimum number of replicates in a cell type to retain it.
#'   Defaults to \code{2}.
#' @param min_features the minimum number of counts for a gene to retain it.
#'   Defaults to \code{0}   
#' @param de_family the differential expression family to use. Available options
#' are:
#' \itemize{
#' \item{"singlecell"}: Uses traditionally methods implemented by Seurat to 
#' test for differentially expressed genes. These methods do not take biological
#' replicate into account. For single cell methods there are \code{six} options
#' for \code{de_method} that can be used, while no input for 
#' \code{de_test} is required:
#' \itemize{
#' \item{"wilcox"}: Wilcoxon Rank-Sum test. The default.
#' \item{"bimod"}: Likelihood ratio test
#' \item{"t"}: Student's t-test
#' \item{"negbinom"}: Negative binomial linear model
#' \item{"LR"}: Logistic regression
#' \item{"MAST"}: MAST (requires installation of the \code{MAST} package).
#' }
#' 
#' \item{"pseudobulk"}: These methods first convert the single-cell expression
#' matrix to a so-called 'pseudobulk' matrix by summing counts for each gene
#' within biological replicates, and then performing differential expression 
#' using bulk RNA-seq methods. For pseudobulk methods there are \code{six} 
#' different methods that can be accessed by combinations of \code{de_method}
#' and \code{de_type}. First specify \code{de_method} as one of the following:
#' \itemize{
#' \item{"edgeR"}: The edgeR method according to
#' Robinson et al, Bioinformatics, 2010. For this method please specify
#' \code{de_type} as either \code{"LRT"} or \code{"QLF"} as the null hypothesis
#' testing approach. See http://www.bioconductor.org/packages/release/bioc/html/edgeR.html
#' for further information. The default.
#' \item{"DESeq2"}: The DESeq2 method according to 
#' Love et al, Genome Biology, 2014. For this method please specify
#' \code{de_type} as either \code{"LRT"} or \code{"Wald"} as the null hypothesis
#' testing approach. See https://bioconductor.org/packages/release/bioc/html/DESeq2.html
#' for further information.
#' \item{"limma"}: The limma method according to
#' Ritchie et al, Nucleic Acids Research, 2015. For this method please specify
#' \code{de_type} as either \code{"voom"} or \code{"trend"} as the precise
#' normalization and null hypothesis testing approach within limma. See
#' https://bioconductor.org/packages/release/bioc/html/limma.html for 
#' further information.
#' }
#' \item{"mixedmodel"}: Mixed model methods also take biological replicate into
#' account by modelling it as a random effect. Please note that these methods
#' are generally extremely computationally intensive. For mixed model methods 
#' there are \code{ten} different methods that can be accessed by 
#' combinations of \code{de_method} and \code{de_type}. 
#' First specify \code{de_method} as one of the following:
#' \itemize{
#' \item{"negbinom"}: Negative binomial generalized linear mixed model. 
#' The default.
#' \item{"linear"}: Linear mixed model.
#' \item{"poisson"}: Poisson generalized linear mixed model.
#' \item{"negbinom_offset"}: Negative binomial generalized linear mixed model
#' with an offset term to account for sequencing depth differences between cells.
#' \item{"poisson_offset"}: Poisson generalized linear mixed model with an 
#' offset term to account for sequencing depth differences between cells.
#' }
#' For each of these options the user has the option to use either a Wald or
#' Likelihood ratio testing method by setting \code{de_type} to \code{"Wald"}
#' or \code{"LRT"}. Default is LRT.
#' }
#' @param de_method the specific differential expression testing method to use.
#' Please see the documentation under \code{de_family} for precise usage options,
#' or see the documentation at https://github.com/neurorestore/Libra. This
#' option will default to \code{wilcox} for \code{singlecell} methods, to
#' \code{edgeR} for \code{pseudobulk} methods, and \code{negbinom} for 
#' \code{mixedmodel} methods.
#' @param de_test the specific mixed model test to use. Please see the 
#' documentation under \code{de_family} for precise usage options,
#' or see the documentation at https://github.com/neurorestore/Libra. This
#' option defaults to \code{NULL} for \code{singlecell} methods, to \code{LRT}
#' for \code{pseudobulk} and \code{mixedmodel} methods. 
#' 
#' @return a data frame containing differential expression results with the 
#' following columns:
#' \itemize{
#' \item{"cell type"}: The cell type DE tests were run on. By default Libra
#' will run DE on all cell types present in the original meta data.
#' \item{"gene"}: The gene being tested.
#' \item{"avg_logFC"}: The average log fold change between conditions.
#' \item{"p_val"}: The p-value resulting from the null hypothesis test.
#' \item{"p_val_val"}: The false discovery rate
#' \item{"de_family"}: The differential expression method family.
#' \item{"de_method"}: The precise differential expression method.
#' \item{"de_type"}: The differential expression method statistical testing type.
#' }
#' 
#' @importFrom magrittr  %<>%
#' @importFrom forcats fct_recode
#' @importFrom dplyr group_by mutate select ungroup arrange
#' @export
#' 
run_de = function(input, 
                  meta = NULL, 
                  replicate_col = 'replicate',
                  cell_type_col = 'cell_type',
                  label_col = 'label',
                  min_cells = 3,
                  min_reps = 2,
                  min_features = 0,
                  de_family = 'pseudobulk',
                  de_method = 'edgeR',
                  de_type = 'LRT') {
  # run differential expression
  DE = switch(de_family,
              pseudobulk = pseudobulk_de(
                input = input, 
                meta = meta, 
                replicate_col = replicate_col,
                cell_type_col = cell_type_col,
                label_col = label_col,
                min_cells = min_cells,
                min_reps = min_reps,
                min_features = min_features,
                de_family = 'pseudobulk',
                de_method = de_method,
                de_type = de_type
              ),
              mixedmodel = mixedmodel_de(
                input = input, 
                meta = meta, 
                replicate_col = replicate_col,
                cell_type_col = cell_type_col,
                label_col = label_col,
                min_features = min_features,
                de_family = 'mixedmodel',
                de_method = de_method,
                de_type = de_type
              ),
              singlecell = singlecell_de(
                input = input, 
                meta = meta,
                cell_type_col = cell_type_col,
                label_col = label_col,
                min_features = min_features,
                de_method = de_method
              )
  )
  
  # clean up the output
  suppressWarnings(
    colnames(DE) %<>%
    fct_recode('p_val' = 'p.value',  ## DESeq2
               'p_val' = 'pvalue',  ## DESeq2
               'p_val' = 'p.value',  ## t/wilcox
               'p_val' = 'P.Value',  ## limma
               'p_val' = 'PValue'  , ## edgeR
               'p_val_adj' = 'padj', ## DESeq2/t/wilcox
               'p_val_adj' = 'adj.P.Val',      ## limma
               'p_val_adj' = 'FDR',            ## edgeER
               'avg_logFC' = 'log2FoldChange', ## DESEeq2
               'avg_logFC' = 'logFC' ## limma/edgeR
    )
    ) %>%
    as.character()
  
  DE %<>% 
    # calculate adjusted p values
    group_by(cell_type) %>%
    mutate(p_val_adj = p.adjust(p_val, method = 'BH')) %>%
    # make sure gene is a character not a factor
    mutate(gene = as.character(gene)) %>%
    dplyr::select(cell_type, 
                  gene, 
                  avg_logFC, 
                  p_val,
                  p_val_adj,
                  de_family, 
                  de_method, 
                  de_type
    ) %>%
    ungroup() %>%
    arrange(cell_type, gene)
}
  
```
