## Supplementary Software 1 for "Confronting false discoveries in single-cell differential expression": singlecell_de.R

```
#' Run single-cell differential expression
#' 
#' Run differential expression using traditional single-cell methods
#' Note that this is effectively a wrapper of the FindMarkers function in Seurat.
#' 
#' @param input a single-cell matrix to be converted, with features (genes) in rows
#'   and cells in columns. Alternatively, a \code{Seurat}, \code{monocle3}, or 
#'   or \code{SingleCellExperiment} object can be directly input.
#' @param meta the accompanying meta data whereby the rownames match the column
#'   names of \code{input}. 
#' @param cell_type_col the vector in \code{meta} containing the cell type 
#'   information. Defaults to \code{cell_type}.
#' @param label_col the vector in \code{meta} containing the experimental
#'   label. Defaults to \code{label}. 
#' @param min_cells the minimum number of cells in a cell type to retain it.
#'   Defaults to \code{3}.
#' @param min_features the minimum number of counts for a gene to retain it.
#'   Defaults to \code{0}   
#' @param de_method the mixed model type to use. Defaults to wilcox.
#' @return a data frame containing differential expression results.
#'  
#' @importFrom magrittr extract set_rownames %<>%
#' @importFrom Matrix rowSums
#' @importFrom dplyr %>% mutate bind_rows
#' @importFrom tibble rownames_to_column 
#' @import Seurat
#' 

singlecell_de = function(
  input,
  meta = NULL,
  cell_type_col = 'cell_type',
  label_col = 'label',
  de_method = 'wilcox',
  min_cells = 3,
  min_features = 0
) {
  
  # check the arguments
  if (!de_method %in% c("wilcox", "bimod", "t", "negbinom", "poisson", "LR",
                        "MAST"))
    stop("Please select one of: wilcox, bimod, t, negbinom, poissoin, LR,
         or MAST as the de_method")
  
  # first, make sure inputs are correct
  inputs = check_inputs(
    input, 
    meta,
    replicate_col = NULL,
    cell_type_col = cell_type_col,
    label_col = label_col
    )
  expr = inputs$expr
  meta = inputs$meta
  
  # define labels, reverse to avoid logFC conflicts
  label1 = unique(meta$label)[2]
  label2 = unique(meta$label)[1]
  
  # get cell types
  cell_types = unique(meta$cell_type)
  
  # create a seurat object
  rownames(meta) = colnames(expr)
  sc = CreateSeuratObject(expr, meta.data = meta)
  
  # make sure idents are set right in the Seurat object
  Idents(sc) = sc$cell_type
  
  # check if integer or already normalized, normalize if needed
  mat = GetAssayData(sc, slot = 'counts')
  if ((sum(mat %% 1 == 0) == length(mat)) == T) {
    sc %<>% NormalizeData()
  } else {
    sc[['RNA']]@data = mat
  }

  # run single cell DE using Seurat
  DE = list()
  for (cell_type_idx in seq_along(cell_types)) {
    cell_type = cell_types[cell_type_idx]
    message("[", cell_type_idx, "/", length(cell_types),
            "] working on cell type: ", cell_type, " ...")
    
    # check to make sure there are enough cells
    n_cells = table(meta$cell_type, meta$label)
    if (min(n_cells[cell_type, ]) < min_cells) {
      message(" .. not enough cells, skipping ...")
      next
    } else {
      tryCatch({
        # subset to the right cell type
        Idents(sc) = sc$cell_type
        sub = sc %>% subset(idents = cell_type)
        # drop genes below threshold
        keep = rowSums(sub) > min_features
        sub = sub[keep,]
        # run DE analysis
        res = FindMarkers(sub, ident.1 = label1, ident.2 = label2,
                              assay = 'RNA', min.pct = -Inf, 
                              min.cells.feature = 0,
                              min.cells.group = min_cells, 
                              logfc.threshold = -Inf,
                              group.by = 'label', 
                              subset.ident = cell_type,
                              test.use = de_method) %>%
          rownames_to_column('gene') %>%
          mutate(
            cell_type = cell_type,
            de_family = 'singlecell',
            de_method = de_method,
            de_type = 'singlecell'
            )
        DE[[cell_type]] = res
      }, error = function(e) message(e))
    }
  }
  DE %<>% bind_rows()
  return(DE)
}
```
