## Supplementary Software 1 for "Confronting false discoveries in single-cell differential expression": to_pseudobulk.R

```
#' Create a pseudobulk matrix
#' 
#' Convert a single-cell expression matrix (i.e., genes by cells)
#' to a pseudobulk matrix by summarizing counts within biological replicates
#' 
#' @param input a single-cell matrix to be converted, with features (genes) in rows
#'   and cells in columns. Alternatively, a \code{Seurat}, \code{monocole3}, or 
#'   or \code{SingleCellExperiment} object can be directly input.
#' @param meta the accompanying meta data whereby the rownames match the column
#'   names of \code{input}.
#' @param replicate_col the vector in \code{meta} containing the replicate 
#'   information. Defaults to \code{replicate}.
#' @param cell_type_col the vector in \code{meta} containing the cell type 
#'   information. Defaults to \code{cell_type}.
#' @param label_col the vector in \code{meta} containing the experimental
#'   label. Defaults to \code{label}. 
#' @param min_cells the minimum number of cells in a cell type to retain it.
#'   Defaults to \code{3}.
#' @param min_reps the minimum number of replicates in a cell type to retain it.
#'   Defaults to \code{2}.
#' @param min_features the minimum number of counts for a gene to retain it.
#'   Defaults to \code{0}   
#' @return a list of pseudobulk matrices, for each cell type.
#'  
#' @importFrom magrittr %<>% extract
#' @importFrom dplyr %>% rename_ count group_by filter pull n_distinct distinct
#'   summarise
#' @importFrom purrr map map_int
#' @importFrom Matrix rowSums colSums
#' @importFrom stats setNames
#' @export
#' 
to_pseudobulk = function(input, 
                         meta = NULL, 
                         replicate_col = 'replicate',
                         cell_type_col = 'cell_type',
                         label_col = 'label',
                         min_cells = 3,
                         min_reps = 2,
                         min_features = 0) {
  # first, make sure inputs are correct
  inputs = Libra:::check_inputs(
    input, 
    meta = meta,
    replicate_col = replicate_col,
    cell_type_col = cell_type_col,
    label_col = label_col)
  expr = inputs$expr
  meta = inputs$meta
  
  # convert to characters
  meta %<>% mutate(replicate = as.character(replicate),
                   cell_type = as.character(cell_type),
                   label = as.character(label))
  
  # keep only cell types with enough cells
  keep = meta %>%
    dplyr::count(cell_type, label) %>%
    group_by(cell_type) %>%
    filter(all(n >= min_cells)) %>%
    pull(cell_type) %>%
    unique()
  
  # process data into gene x replicate x cell_type matrices
  pseudobulks = keep %>%
    map( ~ {
      print(.)
      cell_type = .
      meta0 = meta %>% filter(cell_type == !!cell_type)
      expr0 = expr %>% extract(, meta0$cell_barcode)
      # catch cell types without replicates or conditions
      if (n_distinct(meta0$label) < 2)
        return(NA)
      replicate_counts = distinct(meta0, label, replicate) %>%
        group_by(label) %>%
        summarise(replicates = n_distinct(replicate)) %>%
        pull(replicates)
      if (any(replicate_counts < min_reps))
        return(NA)
      
      # process data into gene X replicate X cell_type matrice
      mm = model.matrix(~ 0 + replicate:label, data = meta0)
      mat_mm = expr0 %*% mm
      keep_genes = rowSums(mat_mm > 0) > min_features
      mat_mm = mat_mm[keep_genes, ] %>% as.data.frame()
      mat_mm %<>% as.data.frame()
      colnames(mat_mm) = gsub("replicate|label", "", colnames(mat_mm))
      # drop empty columns
      keep_samples = colSums(mat_mm) > 0
      mat_mm %<>% extract(, keep_samples)
      return(mat_mm)
    }) %>%
    setNames(keep)
  
  # drop NAs
  pseudobulks %<>% extract(!is.na(.))
  
  # also filter out cell types with no retained genes
  min_dim = map(pseudobulks, as.data.frame) %>% map(nrow)
  pseudobulks %<>% extract(min_dim > 1)
  
  # also filter out types without replicates
  min_repl = map_int(pseudobulks, ~ {
    # make sure we have a data frame a not a vector
    tmp = as.data.frame(.)
    targets = data.frame(group_sample = colnames(tmp)) %>%
      mutate(group = gsub(".*\\:", "", group_sample))
    if (n_distinct(targets$group) == 1)
      return(as.integer(0))
    min(table(targets$group))
  })
  pseudobulks %<>% extract(min_repl >= min_reps)
  return(pseudobulks)
}
```
