## Supplementary Software 1 for "Confronting false discoveries in single-cell differential expression": README.md

# README

Libra is an R package to perform differential expression on single-cell data. Libra implements a total of 22 unique differential expression methods that can all be accessed from one function. These methods encompass traditional single-cell methods as well as methods accounting for biological replicate including pseudobulk and mixed model methods. The code for this package has been largely inspired by the [Seurat](https://satijalab.org/seurat/) and [Muscat](https://github.com/HelenaLC/muscat) packages. Please see the documentation of these packages for further information.

## System requirements

Libra relies on functions from the following R packages:

```
	dplyr (>= 0.8.0),
	purrr (>= 0.3.2),
	tibble (>= 2.1.3),
	magrittr (>= 1.5),
	tester (>= 0.1.7),
	Matrix (>= 1.2-14),
	pbmcapply (>= 1.5.0),
	lmtest (>= 0.9-37),
	tidyselect (>= 0.2.5),
	DESeq2 (>= 0.4.0),
	Seurat (>= 3.1.5),
	blme (>= 1.0-4),
	edgeR (>= 3.28.1),
	glmmTMB (>= 1.0.2.1),
	limma (>= 3.1-3),
	lme4 (>= 1.1-25),
	lmerTest (>= 3.1-3),
	matrixStats (>= 0.57.0),
	methods,
	stats,
	Rdpack (>= 0.7)
```

In addition, the [Seurat](https://satijalab.org/seurat/), [monocle3](https://cole-trapnell-lab.github.io/monocle3/), or [SingleCellExperiment](http://bioconductor.org/packages/release/bioc/html/SingleCellExperiment.html) packages must be installed for Libra to take Seurat, monocle, SingleCellExperiment objects as input, respectively. Methods that require additional packages may also require additional installs (e.g., MAST).

Libra has been tested with R version 3.6.0 and higher.

## Installation

To install Libra, first install the devtools package, if it is not already installed:

```r
> install.packages("devtools")
```

Libra additionally requires the following installations to perform differential expression testing:
```r
> if (!requireNamespace("BiocManager", quietly = TRUE))
    install.packages("BiocManager")

> BiocManager::install(c("edgeR", "DESeq2", "limma"))
```


If Seurat is not installed this will be needed for single-cell methods.
```r
> install.packages("Seurat")
```
Finally, install Libra from GitHub:

```r
> devtools::install_github("neurorestore/Libra")
```

This should take no more than a few minutes.

## Usage

The main function of Libra, `run_de`, takes as input a preprocessed features-by-cells (e.g., genes-by-cells for scRNA-seq) matrix, and a data frame containing metadata associated with each cell, minimally including the cell type annotations, replicates, and sample labels to be predicted.
This means that in order to use Libra, you should have pre-processed your data (e.g., by read alignment and cell type assignment for scRNA-seq) across all experimental conditions.

Libra provides a universal interface to perform differential expression using 22 discrete methods. These methods are summarized as follows:

__Single cell methods__
- Wilcoxon Rank-Sum test
- Likelihood ratio test
- Student's t-test
- Negative binomial linear model
- Logistic regression
- MAST

__Pseudobulk methods__
- edgeR-LRT
- edgeR-QLF
- DESeq2-LRT
- DESeq2-Wald
- limma-trend
- limma-voom

__Mixed model methods__
- Linear mixed model
- Linear mixed model-LRT
- Negative binomial generalized linear mixed model
- Negative binomial generalized linear mixed model-LRT
- Negative binomial generalized linear mixed model with offset
- Negative binomial generalized linear mixed model with offset-LRT
- Poisson generalized linear mixed model
- Poisson generalized linear mixed model-LRT
- Poisson generalized linear mixed model with offset
- Poisson generalized linear mixed model with offset-LRT

By default Libra will use a pseudobulk approach, implementing the `edgeR` package with a likelihood ratio test (LRT) null hypothesis testing framework. Each of the 22 tests can be accessed through three key variables of the `run_de` function: `de_family`, `de_method`, and `de_type`. Their precise access arguments are summarized in the below table.

| Method | de_family | de_method | de_type |
|--------|-----------|-----------|---------|
Wilcoxon Rank-Sum test | singlecell | wilcox | |
Likelihood ratio test | singecell | bimod | |
Student's t-test | singlecell | t | |
Negative binomial linear model | singlecell | negbinom | |
Logistic regression | singlecell | LR | |
MAST | singlecell | MAST | |
edgeR-LRT | pseudobulk | edgeR | LRT
edgeR-QLF | pseudobulk | edgeR | QLF
DESeq2-LRT | pseudobulk | DESeq2 | LRT
DESeq2-Wald | pseudobulk | DESeq2 | Wald
limma-trend | pseudobulk | limma | trend
limma-voom | pseudobulk | limma | voom
Linear mixed model | mixedmodel | linear | Wald
Linear mixed model-LRT | mixedmodel | linear | LRT
Negative binomial generalized linear mixed model | mixedmodel | negbinom | Wald
Negative binomial generalized linear mixed model-LRT | mixedmodel | negbinom | LRT
Negative binomial generalized linear mixed model with offset | mixedmodel | negbinom_offset | Wald
Negative binomial generalized linear mixed model with offset-LRT | mixedmodel | negbinom_offset | LRT
Poisson generalized linear mixed model | mixedmodel | poisson | Wald
Poisson generalized linear mixed model-LRT | mixedmodel | poisson | LRT
Poisson generalized linear mixed model with offset | mixedmodel | poisson_offset | Wald
Poisson generalized linear mixed model with offset-LRT | mixedmodel | poisson_offset | LRT

If batch effects are present in the data, these should be accounted for, e.g., using [Seurat](https://www.sciencedirect.com/science/article/pii/S0092867419305598) or [Harmony](https://www.nature.com/articles/s41592-019-0619-0), to avoid biasing differential expression by technical differences or batch effects.

To run Libra with default parameters on a genes-by-cells scRNA-seq matrix `expr`, and an accompanying data frame `meta`, with `cell_type`, `replicate`, and `label` columns containing cell types, replicates, and experimental conditions, respectively, use the `run_de` function:

```r
> library(Libra)
> DE = run_de(expr, meta = meta)
```

If your columns have different names, you can specify these using the `cell_type_col`, `replicate_col`, and `label_col` arguments:

```r
> DE = run_de(expr, meta = meta, cell_type_col = "cell.type", label_col = "condition")
```

If you would like to store the pseudobulk matrices in a variable, before running differential expression, you can do the following:

```r
> matrices = to_pseudobulk(expr, meta = meta)
```

Libra can also run directly on a Seurat object. For a Seurat object `sc`, with the `sc@meta.data` data frame containing `cell_type` and `label` columns, simply do:

```r
> DE = run_de(sc)
```

The same code can be used if `sc` is a monocle3 or SingleCellExperiment object instead.

## Demonstration

To see Libra in action, load the toy single-cell RNA-seq dataset that is bundled with the Libra package:

```r
> data("hagai_toy")
```

This dataset consists of 600 cells, distributed evenly between six replicates and two conditions.

The `hagai_toy` object is a Seurat object with columns named `cell_type` and `label` in the `meta.data` slot, meaning we can provide it directly as input to Libra:

```r
> head(hagai_toy@meta.data)

                      orig.ident nCount_RNA nFeature_RNA                                       sample replicate label                                  cell_type
2-CGAACATGTATAATGG SeuratProject         18           11 mouse1_lps4_filtered_by_cell_cluster0.txt.gz    mouse1  lps4 bone marrow derived mononuclear phagocytes
2-ATTCTACAGTGGTAGC SeuratProject         23           12 mouse1_lps4_filtered_by_cell_cluster0.txt.gz    mouse1  lps4 bone marrow derived mononuclear phagocytes
2-GCATACACAAACTGTC SeuratProject          3            3 mouse1_lps4_filtered_by_cell_cluster0.txt.gz    mouse1  lps4 bone marrow derived mononuclear phagocytes
2-GAAGCAGAGATGCCAG SeuratProject         14            8 mouse1_lps4_filtered_by_cell_cluster0.txt.gz    mouse1  lps4 bone marrow derived mononuclear phagocytes
2-ATCACGAGTCTAGCGC SeuratProject          3            3 mouse1_lps4_filtered_by_cell_cluster0.txt.gz    mouse1  lps4 bone marrow derived mononuclear phagocytes
2-CTGCCTATCTGTCCGT SeuratProject         28           13 mouse1_lps4_filtered_by_cell_cluster0.txt.gz    mouse1  lps4 bone marrow derived mononuclear phagocytes
```

We run `run_de`, and inspect the differential expression:

```r
> DE = run_de(hagai_toy)
> head(DE)

```

We can also use a different statistical framework, for example `DESeq2`:

```r
> DE = run_de(hagai_toy, de_family = 'pseudobulk', de_method = 'DESeq2', de_type = 'LRT')
> head(DE)
```

Alternatively, we can use a mixed-model approach, which by default will use a negative binomial model structure:

```r
> DE = run_de(hagai_toy, de_family = 'mixedmodel')
> head(DE)
```

However, this can be adjusted using the `de_method` argument:

```r
> DE = run_de(hagai_toy, de_family = 'mixedmodel', de_method = 'linear', de_type = 'LRT')
> head(DE)
```

Running this example on a MacBook should be instantaneous.
However, analyzing >20 real single-cell RNA-seq datasets, we found Libra takes a median of ~5 minutes.
In general, runtime scales close to linearly with the number of cell _types_ and _cells_
If using mixed models, by default, Libra runs on four cores, with each gene analyzed on a different core.
To change the number of cores, use the `n_threads` argument.
For example, running Libra on eight threads:

```r
> DE = run_de(hagai_toy, n_threads = 8)
```

... will run about twice as fast.

## Calculating delta variance
We recently showed that statistical methods for differential expression must account for the intrinsic variability of biological replicates to generate biologically accurate results in single-cell data (Squair et al., 2021, Biorxiv; link). Within the same experimental condition, replicates exhibit inherent differences in gene expression, which reflect both biological and technical factors. We reasoned that failing to account for these differences could lead methods to misattribute the inherent variability between replicates to the effect of the perturbation. To study this possibility, we compare the variance in the expression of each gene in pseudobulks and pseudo-replicates. We call this measure 'delta variance'. Users can use the calculation of delta variance to inform their differential expression results. For example, genes identified as differentially expressed by methods that do not account for biological replicate (i.e., 'singlecell' methods) that have a high delta variance should be treated with caution as they are likely to be false positives. Delta variance can be calculated as follows:

```r
> DV = calculate_delta_variance(hagai_toy)
```

This function will return a list of vectors, one for each cell type, each of which contains the delta variance for the genes present in the input expression matrix.
